# Fundamental equations linking methylation dynamics to maximum lifespan in mammals

**DOI:** 10.1101/2023.05.21.541643

**Authors:** Steve Horvath, Joshua Zhang, Amin Haghani, Ake T. Lu, Zhe Fei

## Abstract

We establish the mathematical foundation that links the rate of change in any molecular biomarker to species lifespan. Specifically, we propose a robust approach that identifies the strong inverse relationship for certain biomarkers using two comprehensive methylation datasets. After examining 54 chromatin states, we found the rates of change of CpG sites in bivalent chromatin states are negatively associated to the lifespans of 90 dog breeds in the first dataset, and the discoveries are further strengthened with 125 mammalian species in the second dataset. Our research leads to three key findings: First, a reciprocal relationship exists between the average rate of methylation change (AROCM) in bivalent promoter regions and maximum lifespan: AROCM ∝ 1/MaxLifespan. Second, the correlation between age and average methylation bears no relation to maximum lifespan, Cor(Methly,Age) ⊥ MaxLifespan. Third, the rate of methylation change in young animals is related to that in old animals: Young animals’ AROCM ∝ Old AROCM. These findings hinge on the chromatin context, as different results emerge when defining AROCM using different chromatin states. Our analytical framework is versatile and readily extendable to a broad range of other molecular assessments. Overall, our study demonstrates that epigenetic aging rates in specific chromatin states exhibit an inverse relationship with maximum lifespan in mammals.

## Introduction

A fundamental question in biology is why closely related species, such as mammals, exhibit significant differences in maximum lifespans (also referred to as maximum longevity or simply longevity). Years of research have outlined the ecological traits associated with maximum lifespan. In short, maximum lifespan is closely linked to an organism’s ability to avoid predation, such as having a large body size, the capacity to fly, or the skill to burrow underground [1, 3, 11, 13, 16, 17, 20]. The rate of living theory suggests that the faster an organism’s metabolism, the shorter its lifespan. This theory was initially proposed by Max Rubner in 1908, following his observation that larger animals lived longer than smaller ones, and that these larger animals had slower metabolisms. Rubner’s rate of living theory, which was once accepted, has now largely been debunked. This shift in perception is due to the application of modern statistical methods which account for the effects of both body size and phylogeny. When these factors are appropriately adjusted for, there is no correlation between metabolic rate and lifespan in mammals or birds [16, 17]. Nonetheless, contemporary adaptations of the original rate of living theory have emerged. Many articles have explored the relationship between the rate of change in various molecular markers and maximum lifespan. Specifically, maximum lifespan has been connected to the rates of change in telomere attrition [5, 12, 26, 49, 50, 53], somatic mutations [7, 9, 36, 43, 48], and cytosine methylation [22, 33, 35].

The strong correlation between maximum lifespan and the rate of change in various factors (telomeres, somatic mutations, methylation) raises the possibility that these relationships might merely be artifacts resulting from the definition of the rate of change. In other words, the strong correlation with maximum lifespan could be a mathematical consequence stemming from the calculation of rates of change per year. To address this concern, we introduce a framework that links the rate of change in a biomarker to maximum lifespan. We show that any biomarker positively correlated with age in multiple species will exhibit an inverse relation with maximum lifespan. Thus considerable care must be taken to avoid biases deriving from the definition of the rate of change.

## Results

We examine the correlation between methylation dynamics throughout the lifespan and maximum lifespan using two datasets from the Mammalian Methylation Consortium [23]. The first dataset encompasses blood methylation data from 90 distinct dog breeds, while the second dataset encompasses many different tissue types from 125 mammalian species. We begin with the simpler dog dataset, which consists of a single tissue, to familiarize the reader with our methodology. Our primary scientific emphasis is on the mammalian lifespan dataset, which we analyze in depth.

### Rate of Change in Methylation in Dog Breeds

Dog breeds exhibit a striking range of lifespans, with some breeds living up to twice as long as others. A prior study involving two dog breeds indicated an inverse relationship between methylation change rate and breed-specific lifespan [33]. Here we examine the connection between methylation change rate and the lifespans of dog breeds using a substantially larger dataset: *N* = 742 distinct blood samples from 90 different dog breeds [28]. For later applications, we will use the more abstract term “stratum” to replace “dog breed”. We first introduce the definition of Average Rate of Change in Methylation (AROCM), which quantifies the rate or gradient of age-related changes in a set of cytosines across samples from a given stratum. Therefore AROCMs depend on the age interval and the set of cytosines. Assuming there are *n* (blood) samples within one stratum (dog breed) and an age range [*ℓ, u*], the methylation matrix is

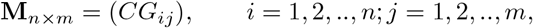

where rows represent (blood) samples and columns represent a set of *m* CpGs (e.g., CpGs located in a specific chromatin state). For the *i*-th sample, the average methylation value *Methyl* is defined as:

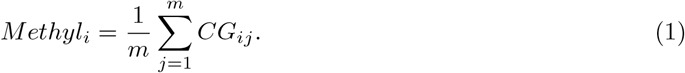

To enable comparisons with other aging biomarkers, we define the scaled methylation value as:

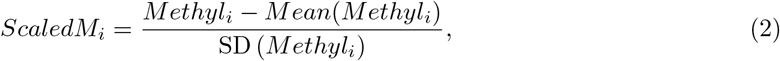

where the Mean and SD are taken over samples *i* = 1, 2, ‥, *n* within each stratum. Now we define the AROCM in one stratum as the coefficient estimate *β*_1_ resulting from the univariate linear regression model:

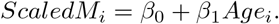

The term “average” in AROCM reflects that *Methyl* was defined as the average value across a specific set of cytosines (equation 1). In our analysis of the dog data, we concentrated on the 552 CpGs situated within the BivProm2+ chromatin state, also known as the Bivalent Promoter 2 state that is associated with the Polycomb Repressive Complex 2 (PRC2). The rationale behind selecting this specific chromatin state will be presented in our subsequent application concerning mammalian maximum lifespan. A comprehensive description of chromatin states can be found in the Methods section. With some detailed derivation (**Methods**), we have

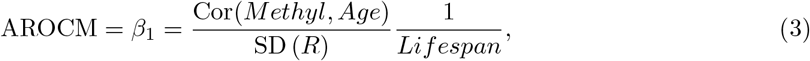

where 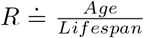 is the relative age in each stratum. Equation (3) reveals that the inverse association between AROCM and maximum lifespan would hold when the first ratio term 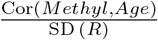 approximates some constant across the strata (Figure 1a,b). This raises the concern that the relationship between the rate of change (AROCM) and 1*/Lifespan* might simply result from selecting CpGs with a positive age correlation. To mitigate this concern, we avoided pre-filtering CpGs based on their age correlations. We chose to define the rate of change in relation to sets of CpGs associated with chromatin states that were defined with respect to histone marks (refer to **Methods**). In practical data applications, inconsistent age ranges and the associated variability in SD (*R*) notably influence the estimation of AROCM and Cor(*Age, Methyl*). The dependence of AROCM and Cor(*Age, Methyl*) on SD (*R*) usually is not of biological interest. Instead, it indicates flaws in sample selection and study design. To mitigate the impact of these sampling imperfections, we introduce adjusted values:

**Figure 1:**
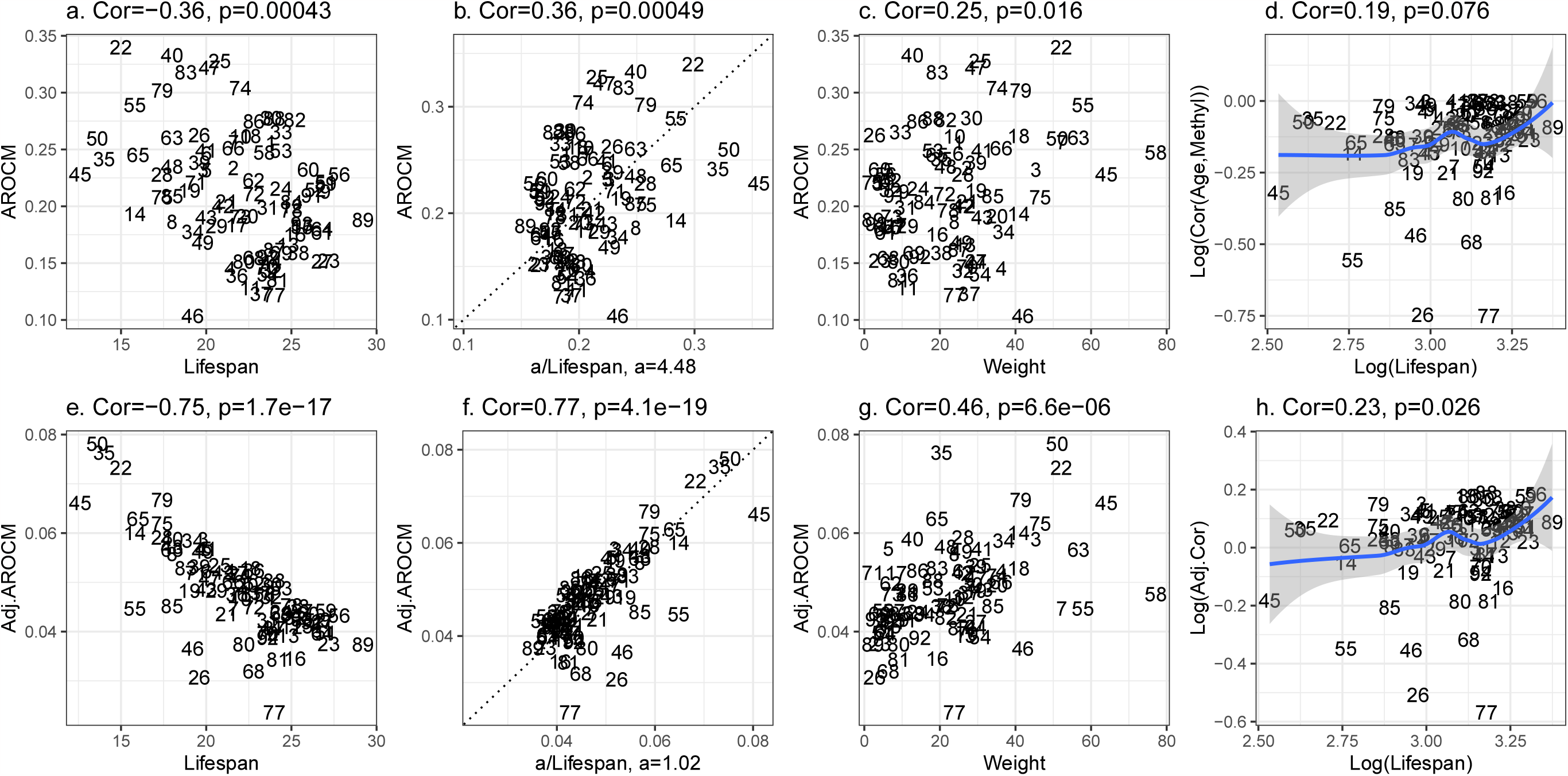
AROCMs in blood samples from *S* = 90 dog breeds. Top panels connect AROCMs to **a**. Lifespan, **b**. 1*/Lifespan*, **c**. adult weight, and **d**. Log(Cor(Age, Methyl)) versus Log(Lifespan); bottom panels tie *Adj*.*AROCM* to the same factors **(e**,**f**,**g)**, and **h**. Log(Adj.Cor) versus Log(Lifespan). Both adjusted and unadjusted AROCMs were computed with *p* = 552 CpGs in bivalent promoter 2 bound by polycomb repressive complex 2 (PRC2) (BivProm2+)[51]. Each integer label corresponds to a different dog breed (Supplementary Table 1).

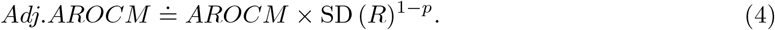

This straightforward adjustment multiplies *AROCM* by a power term of SD (*R*). It offers the additional benefit of establishing a simple relationship with maximum lifespan (**Methods**):

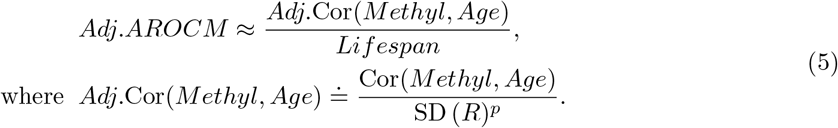

To arrive at a pronounced inverse correlation between Adj.AROCM and Lifespan, equation 5 suggests minimizing the coefficient of variation in *Adj*.Cor(*Methyl, Age*) (see equation 25 in **Methods** for more details). When selecting *p*, we do not take lifespan information into account. However, this criterion for choosing *p* faces a statistical challenge: overfitting, i.e. resulting in an artificially tight inverse relation between the rate of change and lifespan. Therefore, we advise presenting results for both the original and adjusted values of AROCM in practical applications. Since *p* = 0 implies *Adj*.Cor = Cor and no adjustment, smaller *p* indicates a lesser degree of sampling imbalance and therefore smaller adjustment. According to the criterion for choosing the adjustment power, *p* = 0.1 emerges as a suitable choice for our dog methylation data (Supplementary Figure S1).

The low correlation between *Adj*.Cor(*Methyl, Age*) and breed lifespan on a logarithmic scale (r=0.23, p=0.026, Figure 1h) implies strong negative relationship between Adj.AROCM and lifespan on the log scale, as indicated by Proposition 2. This is validated with a correlation coefficient of *r* = − 0.75 (Figure 1e, Supplementary Table 1). We also identify positive correlations between AROCM and average breed weight (Figure 1c,g), which is predictable since breed weight inversely correlates with breed lifespan.

While the adjusted version of AROCM, Adj.AROCM, comes with drawbacks such as the risk of overfitting and reduced interpretability, it bears several benefits. First, it often boosts the data signal (*r* = − 0.75 for the adjusted AROCM versus *r* = − 0.36 for the unadjusted AROCM, Figure 1a,e). Second, it aids in deriving formulas for constants of proportionality (equation 23 in **Methods**). For instance, the constant is 1.02 in equation (23) among dog breeds in our data (Figure 1f). In summary, both the adjusted and unadjusted AROCMs display negative correlations with lifespan when viewed on a logarithmic scale. Yet, the correlation is notably stronger in the adjusted AROCM when compared to its unadjusted counterpart. This indicates that the adjusted version adeptly counteracts the limitations of an imperfect dataset, particularly the confounding influence of varying *SD*(*R*) values.

### Mammalian Methylation Data and Chromatin States

Recent investigations have revealed a connection between the rates of methylation change and the maximum lifespan of mammals [33]. In the present study, we revisit this question utilizing a comprehensive dataset from the Mammalian Methylation Consortium [23, 34]. These data are well suited for comparative aging rate studies, as the mammalian array platform provides a high sequencing depth at CpGs, which are highly conserved across mammalian species [2]. We maintained the same definitions as in our dog dataset, though strata were classified by species and tissue types within species. In order to derive reliable AROCM estimates, we removed outlying species/tissue strata using criteria detailed in **Methods**. The analysis of the AROCM defined with respect to the entire age range [0, *Lifespan*) involves *S* = 229 distinct species/tissue strata. We first calculated AROCMs in all species-tissue strata (equation 3) and aggregated them by species for all 54 chromatin states identified on the mammalian array (Figure 2, Supplementary Table 2). Specifically, the species-level AROCM is defined as the median across tissue types within that species.

**Figure 2:**
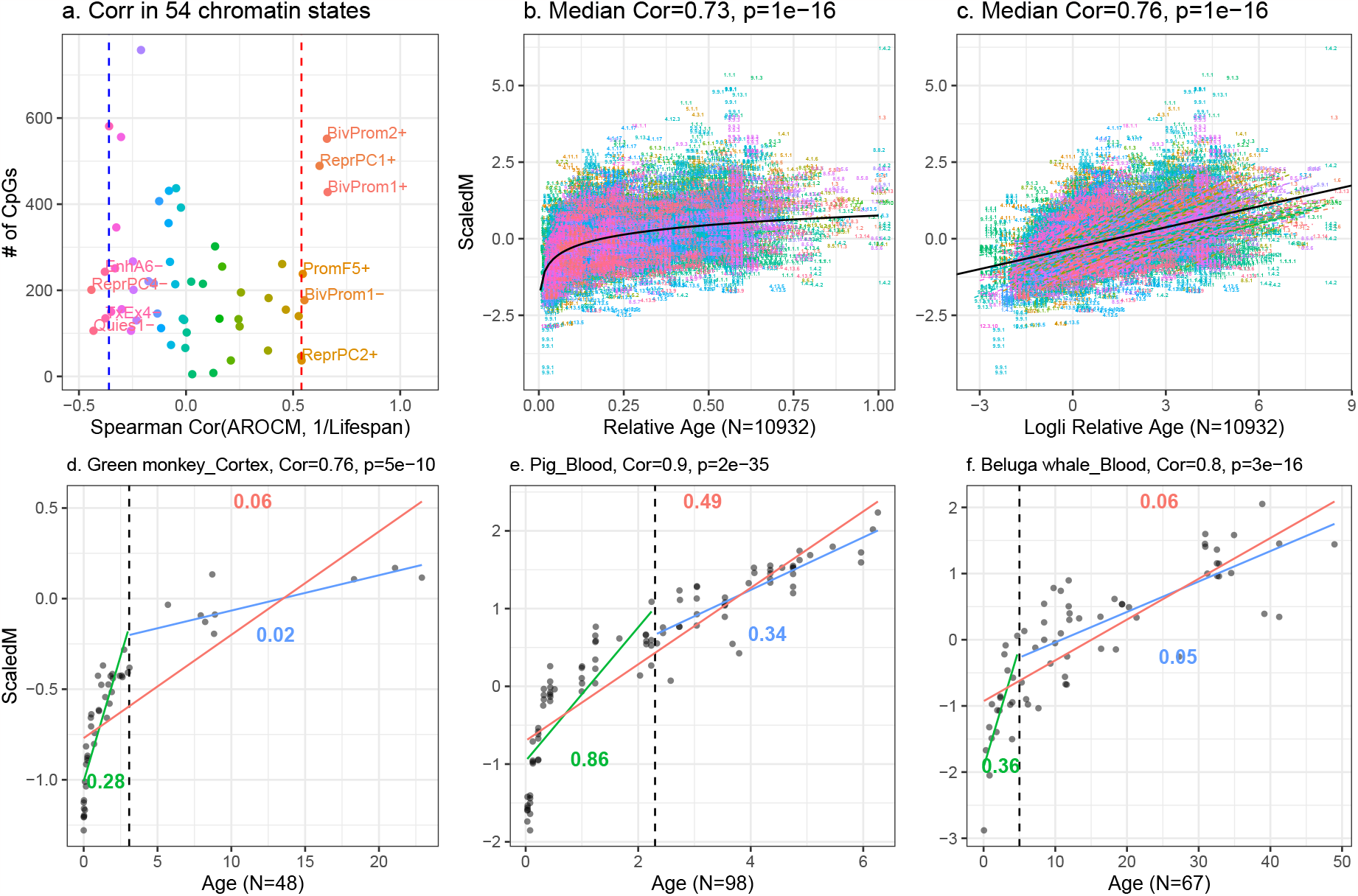
Age versus mean methylation in select chromatin states in mammals. **(a)** Displays the Spearman correlation between the inverse of mammalian maximum lifespan 1*/L* (x-axis) and the AROCM of diverse chromatin states. Each dot represents one of the 54 universal chromatin states [51], with the AROCM defined per Methods. Only chromatin states with Spearman correlation exceeding 0.6 or below −0.3 with the inverse of maximum lifespan are labeled. More details are in Supplementary Table 2. The most pronounced correlation is found for CpGs in bivalent promoter 2 also bound by polycomb repressive complex 2 (BivProm2+). **(b)** Scaled Mean Methylation versus Relative Age across all samples; **(c)** Scaled Mean Methylation against Log-linear Transformed Relative Age in all samples. Black curves mark the overall trend. In panel **(c)**, a linear regression line was fitted per species. **(d-f)** Depicts AROCM calculation for BivProm2+ state in 3 species-tissue strata. Colored line segments/numbers in each panel represent: Young AROCM for age interval [0, 0.1*L*) in green; Old AROCM for age interval (0.1*L, L*) in blue; AROCM for age interval [0, *L*) in red.The sample size for each panel is reported in the x-axis label.

Given that AROCM estimates may exhibit a non-linear relationship with the inverse of maximum lifespan (Figure 2b), we calculated the Spearman correlation (as opposed to the Pearson correlation) between AROCM and 1*/L*, the lifespan. The Spearman correlation coefficients of all chromatin states varied from −0.44 (ReprPC4-) to 0.66 (BivProm1+, BivProm2+) (Figure 2a, Supplementary Table 2). First, we will discuss chromatin states such as TxEx4-, ReprPC4-, and Quies1-. These states typically exhibit high methylation values [23] and their AROCM estimates frequently result in negative values. Such chromatin states, with frequently negative AROCM estimates, typically exhibit a negative correlation with 1*/Lifespan*, as indicated by the Spearman correlation in Figure 2a. For these states, lower negative AROCM values correspond to shorter lifespans. Essentially, species experiencing rapid age-related methylation loss in TxEx4-, ReprPC4-, and Quies1-generally have shorter lifespans. However, the lifespan correlations for chromatin states with negative AROCM values are not as pronounced as those for states with positive AROCM values. We will primarily focus on chromatin states that typically exhibit low methylation values and their AROCM estimates frequently result in positive values. Specifically, the 552 CpGs in the BivProm2+ chromatin state show the strongest positive correlation with 1*/Lifespan*. This is why we used this chromatin state in our prior application to dog breeds. It’s worth noting that similar results can be observed in other chromatin states with low methylation values, such as BivProm1+ and ReprPC1+.

### Non-linear relationship across the life course

In various species-tissue strata, a non-linear relationship exists between age and scaled mean methylation *ScaledM* (equation 2), evident in species like the green monkey, pig, and beluga whale (Figure 2d-f). A similar non-linear pattern is observed for relative age *R* (Figure 3). Our Methods section details a parametric model for this relationship, as described in equations 27 and 28. This non-linearity poses a mathematical challenge in estimating AROCM. However, this can be addressed by dividing the age range into segments where linear relationships are appropriate (Figure 2d-f). Roughly linear relationships can be observed when focusing on either young animals (defined by *R <* 0.1) or older animals (*R* ≥ 0.1). For instance, in humans, we designated the young stratum by *R <* 0.1, corresponding to *Age <* 12.2 years, while the older stratum was defined as *Age* ≥ 12.2. Using the definitions of ‘Young’ and ‘Old’, we computed the AROCM for three specific chromatin states: BivProm2+, BivProm1+, and ReprPC1+. For all three states, we observed that two main results: i) AROCM values are higher in young animals than in older ones; ii) a pronounced positive correlation exists between AROCM values in young and old animals, with a Pearson correlation coefficient of *r >* 0.78 (see Figure 4). While both biological and mathematical factors (detailed in the Methods section) account for the higher AROCM in young animals, the robust positive correlation (*r >* 0.78) stands out. This insight was made possible due to the extensive sample size provided by our Mammalian Methylation Consortium. To address any concerns regarding the 0.1 threshold for *R*, we performed comprehensive sensitivity analyses using thresholds from 0.2 to 0.9, confirming the robustness of our findings (see Supplementary Table 3).

**Figure 3:**
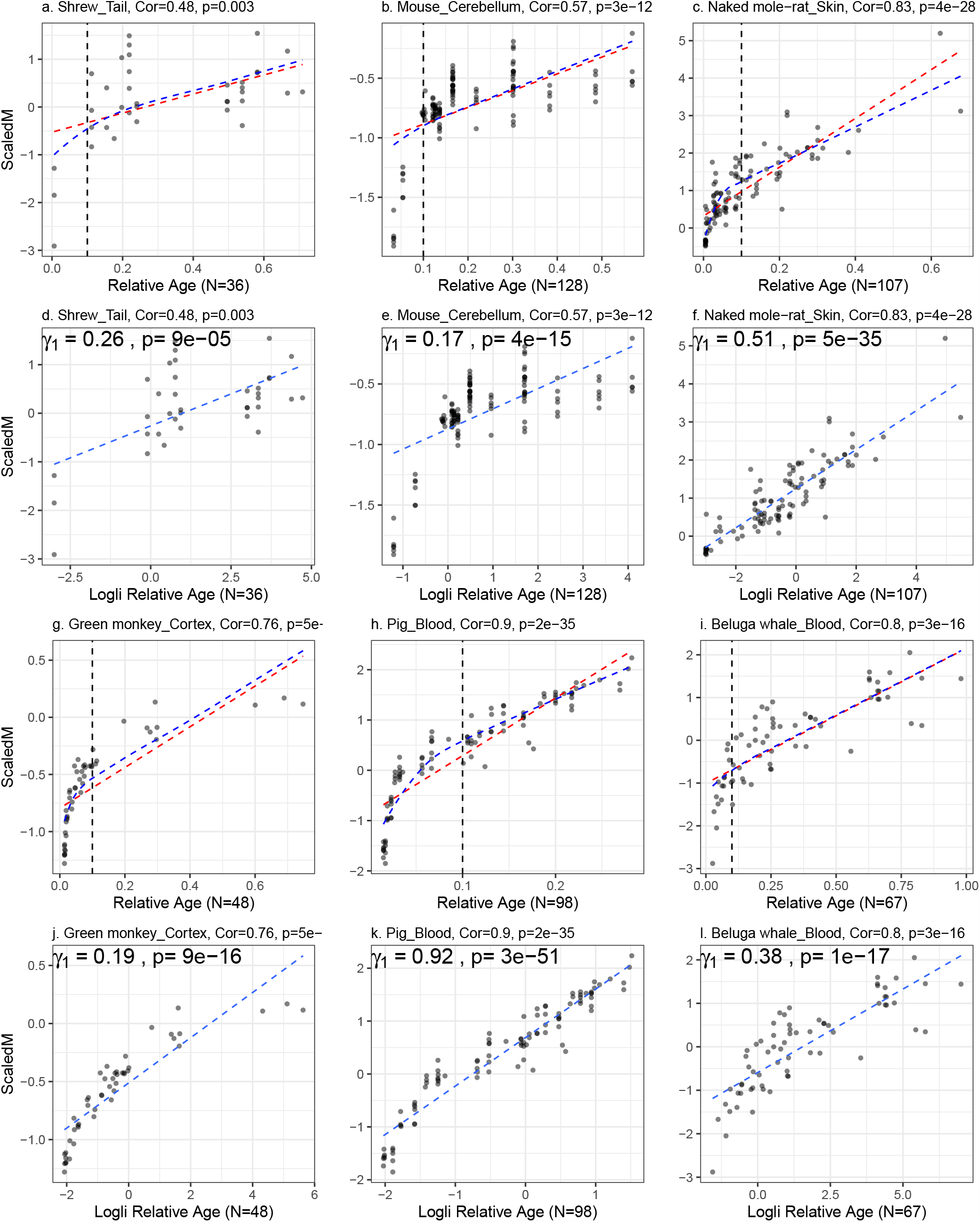
Scaled Mean Methylation against Relative Age or Loglinear Transformed Relative Age in 6 Additional Species Tissue Strata. The correlations increase and associations become more linear post transformation. **(a-c, g-i)** Scaled Mean Methylation versus Relative Age, red dashed line represents linear fit to relative age, blue dashed curve represents the inverse transformation of the linear fit in (d-f, j-l); **(d-f, j-l)** Scaled Mean Methylation versus Loglinear transformed relative age, where *γ*_1_ is the AROCM defined in equation (27), with *ScaledM*_*i*_ = *γ*_0_ +*γ*_1_*T*_*i*_. The sample size for each panel is reported in the x-axis label.

**Figure 4:**
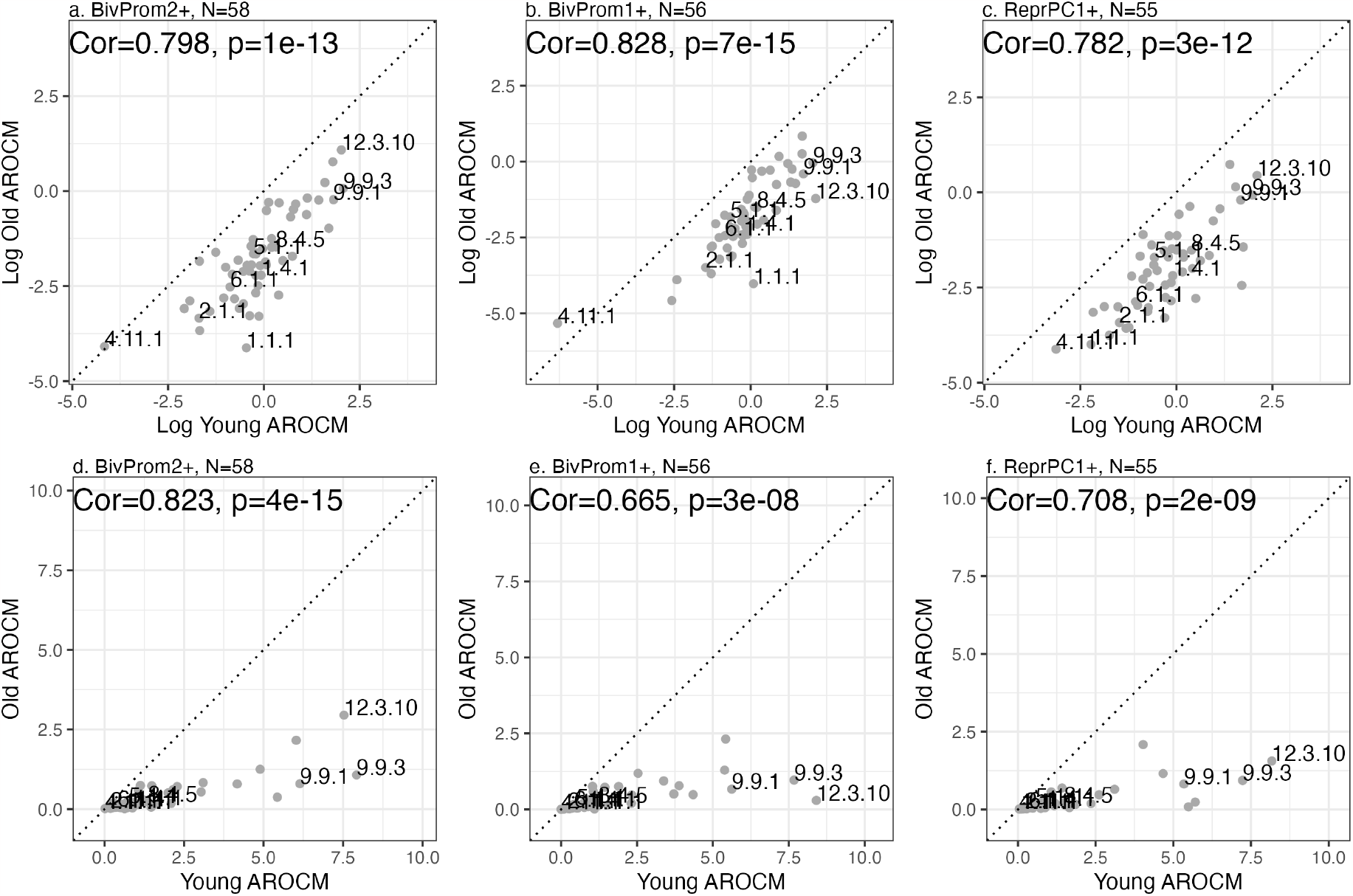
AROCM in young animals versus that in old animals. **(a**,**b**,**c)** Natural log transformed values of AROCM in young animals versus AROCM calculated in old animals. Mean methylation was defined with respect to CpGs located in chromatin state **(a)** BivProm2+, **(b)** BivProm1+, **(c)** ReprPC1+. **(d**,**e**,**f)** Corresponding plots on the original scale, i.e. without log transformation. Each dot corresponds to a different species. Median aggregation was used to combine AROCM results from different tissues by species. X-axis: AROCM for young samples (Relative Age *R <* 0.1). Y-axis: AROCM for old samples (*R >*= 0.1). Samples are labelled by mammalian species numbers. Each panel reports the Pearson correlation value Cor and the corresponding p-value.

### Adjusted AROCM approximates the inverse of maximum lifespan

Given that the coefficient of variation for *Adj*.*AROCM* is substantially lower than that for *Lifespan*, our mathematical framework predicts an inverse correlation between *Adj*.*AROCM* and *Lifespan*, our mathematical formalist predicts an inverse relationship between *Adj*.*AROCM* and *Lifespan*, as indicated by equation 5 and Proposition 2. Indeed, both the AROCM (*r* = − 0.85) and the adjusted AROCM (*r* = − 0.92) exhibit strong negative correlations with maximum lifespan on the log scale (Figure 5a,c). The difference between the two measures is more pronounced when looking on the original scale (no log transformation): the adjusted AROCM leads to a high correlation with *a/Lifespan* (*r* = 0.96) compared to that for the unadjusted AROCM (*r* = 0.72, Figure 5b,d). A subtle benefit of the adjusted correlation *Adj*.Cor(*Methyl, Age*) is that its mean value approaches 1.0 across species (mean = 1.11, Supplementary Figure S2i) since this results in a simple memorable formula: the adjusted AROCM is roughly equal to 1*/Lifespan* because *a* ≈ 1 (equation 5, Figure 5d). Strong negative correlations between lifespan can be further observed when AROCM is defined with respect to young animals only (*R <* 0.1) or old animals only (*R* ≥ 0.1, Supplementary Figure S3). Since maximum lifespan is positively correlated with average age of sexual maturity and gestation time, we find that both variables correlate strongly with AROCM as well (Supplementary Figure S4).

**Figure 5:**
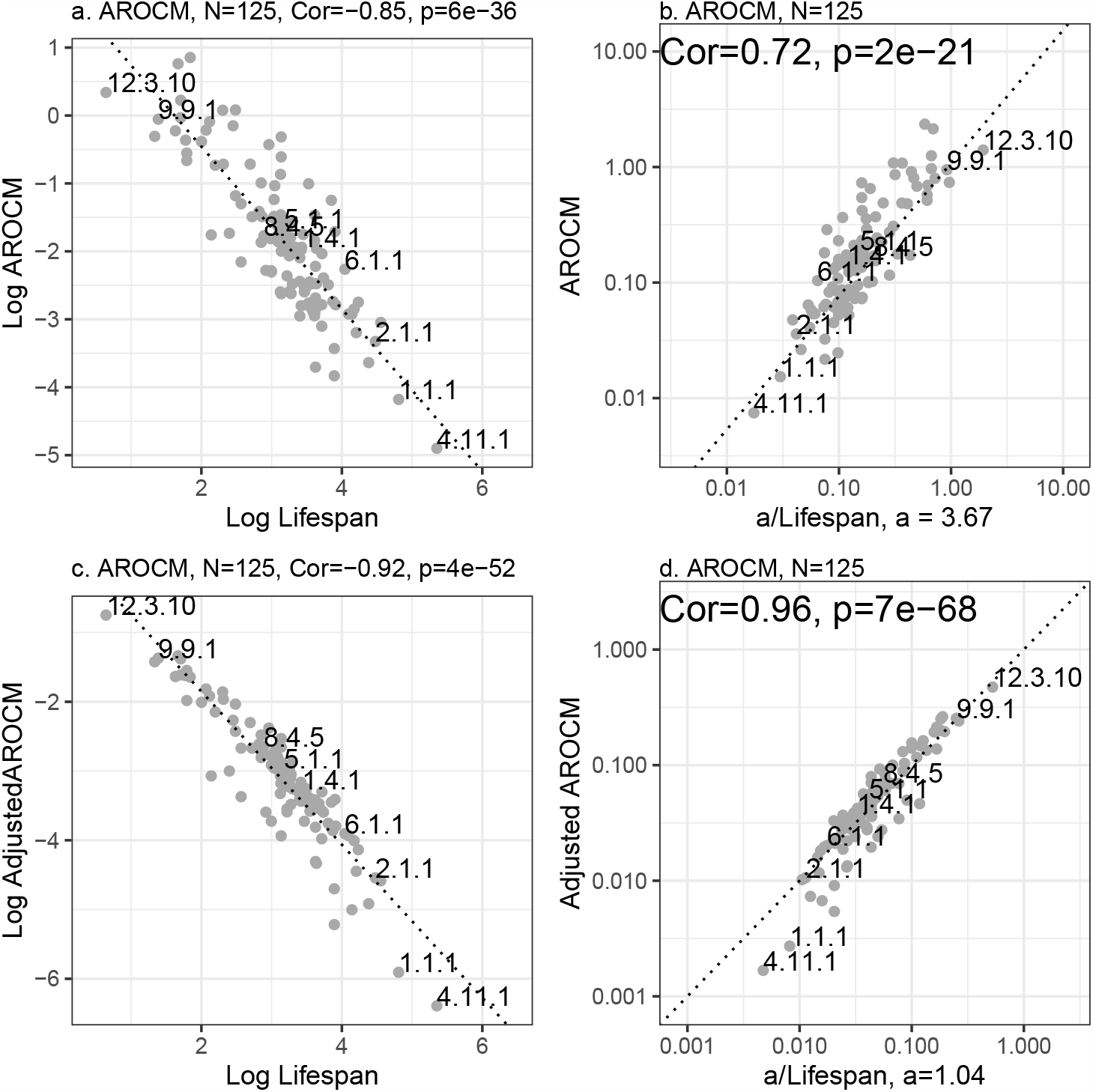
AROCM Relationship with Maximum Mammalian Lifespan. **(a)** Log AROCM versus Log Lifespan; **(b)** AROCM against *a/Lifespan*; **(c)** Log Adj.AROCM versus Log Lifespan (equation 19); **(d)** Adj.AROCM against *a/Lifespan* (equation 23). Figures represent species-level results (median across tissues) with 125 species. Pearson correlations are reported for each panel. Dotted lines in **(b**,**d)** denote *x* = *y*. Supplementary Figure S3 contains results for young and old AROCM.

### Methylation-age correlation does not relate to lifespan

The correlation between *Methyl* and *Age*, denoted as Cor(*Methyl, Age*), assumes similar values for long and short lived species, e.g. Median Cor(*Methyl, Age*) =0.51 for humans, 0.58 for humpback whale, 0.62 for Asian elephant, 0.66 for mouse, 0.67 for brown rat, 0.69 for prairie vole. To delve deeper into the relationship between Cor(*Methyl, Age*) and maximum lifespan, we grouped each species into one of five groups based on their maximum lifespan: less than 10 years, 10-19 years, 20-24 years, 25-39 years, or over 40 years. These groupings were designed to ensure a balanced number of species per group. Our analysis revealed no discernible trend between either Cor(*Methyl, Age*) or *Adj*.Cor and lifespan group (Figure 6). Similarly, we find no relationship between lifespan and Cor(*Methyl, Age*) when the latter is defined with respect to young or old animals (**Supplementary Methods**). The analysis mentioned above overlooks a technical challenge. In our dataset, Cor(*Methyl, Age*) shows a correlation with the standard deviation of relative age, with a Pearson correlation of *r* = 0.23 in all species tissue strata and *r* = 0.27 across species (Supplementary Figure S2d,e). This correlation is not ideal, as it stems from an imbalanced and imperfect data sampling. Ideally, with all species sampled to have the same distribution in relative age, SD (*R*) would remain constant, making Cor(*Methyl, Age*) and SD (*R*) independent (**Supplementary Methods**). The variability in SD (*R*) led to the introduction of the adjusted correlation, *Adj*.Cor (equation 4). Using an adjustment power of *p* = 0.25 (Supplementary Figure S1b), *Adj*.Cor is found to have non-significant correlation with SD (*R*) (Supplementary Figure S2g,h).

**Figure 6:**
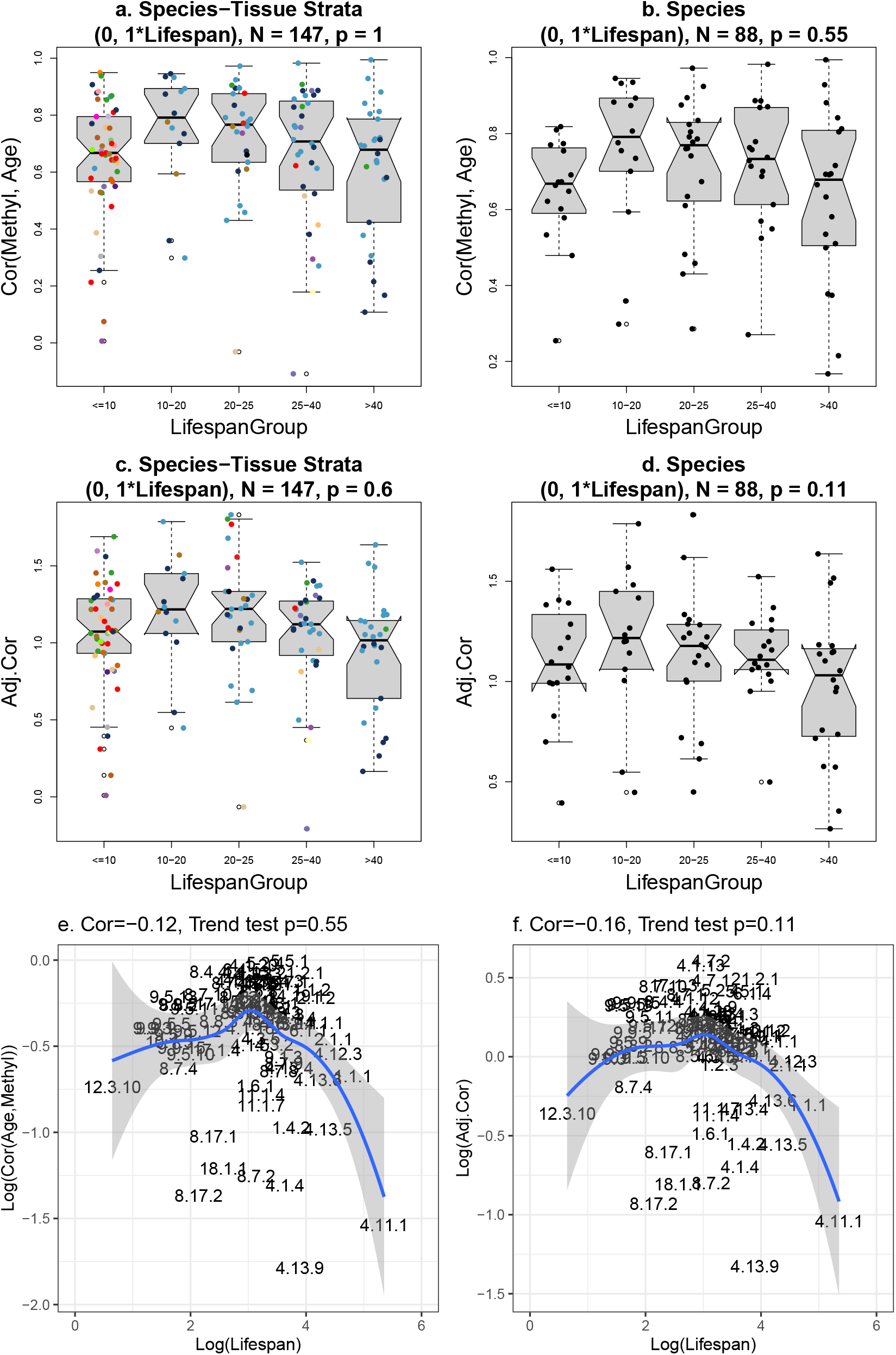
Lifespan group analysis of age correlations. Cor(*Methyl, Age*) and 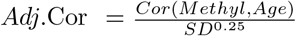 versus groups of species defined by maximum lifespan. **(a, b)** Non-significant associations between *Cor*(*Methyl, RelAge*) and lifespan groups. **(c, d)**: Adjusted Cor (equation 17) by Lifespan groups. **(e, f)**: Log of Cor(Methyl,Age) and Adj.Cor versus Log of Lifespan. P-values are based on the Mann-Kendall Trend Test. These results illustrate condition (C1) in Methods. For each box, the dots correspond to individual strata, the center line is the median; the box limits are upper and lower quartiles; the whiskers are 1.5 times interquartile range.

### Adult weight does not confound the relationship between AROCM and lifespan

Body size (e.g. measured using average adult body mass or adult weight) is a major determinant of maximum lifespan—with larger animals living, on average, longer than smaller ones [8, 10, 16, 31, 40, 42, 45]. Adult weight is often a confounding factor in comparative studies of maximum lifespan [25, 40, 44]. To address this concern, we carried out several analyses to demonstrate that adult weight does not explain the observed relationships between AROCM and maximum lifespan. First, we demonstrate that the correlation between log(AROCM) and log(Weight) is much weaker than that observed for maximum lifespan (*r* = − 0.66 for average weight, Supplementary Figure S5a, versus *r* =− 0.85 for maximum lifespan, Figure 5a). Second, we fit the following multivariate regression models between the unadjusted AROCM, maximum lifespan and adult weight (Table 1),

**Table 1:** Regression models involving maximum lifespan. We considered animals across the entire age range. We used mean methylation in BivProm2+ for all species-tissue strata (*N* = 229).

| Model | Covariate | Estimate | Std. Error | t value | Pr(> \|t\|) |
| --- | --- | --- | --- | --- | --- |
| M1: $\log(Lifespan) \sim$ | (Intercept) | 1.54 | 0.071 | 21.7 | 5.40E-52 |
| | $\log(AROCM)$ | -0.44 | 0.026 | -16.92 | 1.80E-35 |
| | $\log(AverageWeight)$ | 0.09 | 0.01 | 9.23 | 5.10E-18 |
| M2: $\log(AROCM) \sim$ | (Intercept) | 1.92 | 0.18 | 10.94 | 9.80E-17 |
| | $\log(Lifespan)$ | -1.3 | 0.077 | -16.92 | 1.80E-35 |
| | $\log(AverageWeight)$ | 0.015 | 0.02 | 0.76 | 0.45 |
| M3: $\log(AROCM) \sim$ | (Intercept) | 1.7 | 0.19 | 9 | 1.20E-16 |
| | $\log(Lifespan)$ | -1.2 | 0.079 | -15 | 1.80E-35 |
| | $\log(AverageWeight)$ | 0.015 | 0.02 | 0.78 | 0.43 |
|  | Brain | -0.26 | 0.19 | -1.4 | 0.16 |
| M4: $\log(AROCM) \sim$ | (Intercept) | 1.79 | 0.22 | 8.24 | 1.94e-14 |
| | $\log(Lifespan)$ | -1.28 | 0.08 | -16.06 | 2e-16 |
| | $\log(AverageWeight)$ | 0.03 | 0.02 | 1.22 | 0.23 |
|  | Brain | 0.03 | 0.21 | 0.16 | 0.87 |
|  | Blood | 0.16 | 0.15 | 1.02 | 0.31 |
|  | Skin | 0.11 | 0.16 | 0.71 | 0.48 |
|  | Liver | 0.50 | 0.23 | 2.13 | 0.03 |
|  | Muscle | -0.52 | 0.37 | -1.42 | 0.16 |
|  | Tail | 0.19 | 0.33 | 0.59 | 0.56 |

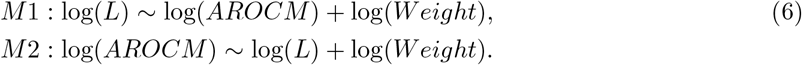

When lifespan is regressed on AROCM and weight on the log scale, both AROCM and weight are highly significant (model M1, Table 1). When AROCM is regressed on lifespan and weight on the log scale, lifespan remains highly significant while weight is not (model M2, Table 1). These results demonstrate that adult weight has only a weak effect on the relationship between lifespan and AROCM.

### Effect of tissue type on the relationship between AROCM and lifespan

Most proliferating tissue types have similar values of Cor(*Methyl, Age*) but slightly lower correlations can be observed for non-proliferating tissues such as the cerebellum, cerebral cortex, skeletal muscle, and heart when looking at all samples irrespective of age (Supplementary Figure S6). To carry out a formal analysis of the effect of tissue type, we added indicator variables for tissue types as covariates to the multivariate regression models that explored the relationship between AROCM and lifespan:

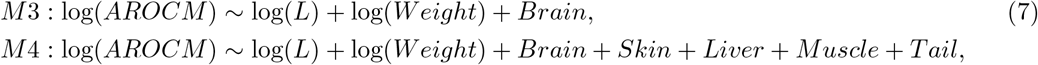

where model M4 added indicator variables for several tissue types. Both M3 and M4 allow insights into the potential tissue differences in AROCM estimates, as detailed estimates can be found in Table 1. After including log(*L*) as the covariate, only liver tissue had a significant (positive) association with log(*AROCM*), (*p* = 0.03).

### Phylogenetic effects

The shared genetic, behavioral, and ecological traits among related species, inherited from common ancestors, can create non-random associations or ‘phylogenetic signals’. Ignoring these phylogenetic relationships can lead to false correlations or overlook real ones [19, 25, 40]. Upon adjusting for phylogenetic relationships, the previously mentioned associations persist, albeit with weaker correlation coefficients. In the case of the dog methylation data, we observed substantial positive associations in the phylogenetically independent contrasts (PICs [19]) of AROCM and breed lifespan (refer to Supplementary Figure S7). Here, we recorded a correlation of − 0.26 (*p* = 0.018) for AROCM, and a correlation of − 0.64 (*p* = 7.3 *×*10^−11^) for Adjusted AROCM.

In the mammalian data, we again found significant positive associations in PICs of AROCM and maximum lifespan (Supplementary Figure S8). Correlations for this dataset were *r* = − 0.2 (*p* = 0.03) for AROCM, − 0.35 (*p* = .00013) for Adjusted AROCM, and − 0.23 (*p* = .022) for Adjusted Old AROCM. Collectively, these findings indicate a connection between the rate of methylation change in bivalent promoter regions and maximum lifespan, which holds even after phylogeny adjustment.

## Discussion

### The rate of change isn’t always inversely proportional to lifespan

Prior studies have linked the rate of change in methylation to maximum lifespan in smaller samples of mammalian species [22, 33, 54], and have suggested a strong positive correlation between methylation rate changes and the inverse of mammalian lifespan. But this is not always the case. By leveraging the large data set from our Mammalian Methylation Consortium and a careful mathematical framework we demonstrate that the strong positive correlation between AROCM and 1/lifespan are only found in certain chromatin states such as bivalent promoters (Figure 2a). In other chromatin states, the correlation between AROCM and 1/lifespan is either not present or even reversed (Supplementary Figure S9). Future studies might delve deeper into which chromatin regions result in the opposite interpretation, where a rapid rate of change correlates with an *increased* maximum lifespan. Our research initiated this exploration by pinpointing chromatin states where the rate of change negatively correlates with 1*/Lifespan*. One intriguing aspect is understanding why chromatin states like TxEx4-, ReprPC4-, and Quies1, which negatively correlate with 1/Lifespan, exhibit a more subdued p-value association compared to their positively correlated counterparts. This might be attributed to the varied age-related methylation loss patterns across different tissue types. For example, the TxEx4 high-expression transcription state indicates age-related methylation loss in non-proliferative tissues [34]. Yet, in proliferative tissues such as blood or skin, TxEx4 has diminished enrichment of age-related cytosines undergoing methylation loss [34]. Further, the correlation between mammalian lifespan and AROCM is not significant when employing all 8970 CpGs that correspond to both eutherians and marsupials (Supplementary Figure S10). These findings collectively underscore the significance of selecting an appropriate chromatin context for defining the AROCM.

### Chromatin state BivProm2 is special

Bivalent chromatin states generally exhibit low methylation levels in most tissues and species and are bound by polycomb repressive complex 2 (PRC2) [34, 37, 47]. Many previous articles, including our pan-mammalian aging studies, have shown target sites of PRC2 gain methylation with age in most species and tissues [34, 37, 41, 47]. We proposed a simple and fundamental equation that links the adjusted rate of change in bivalent chromatin regions to the inverse of maximum lifespan (equation 5). Note that equation (5) does not involve hidden parameters. Our study strongly suggests that careful definition and measurement of both *Adj*.*AROCM* ^(*s*)^ and *L*^(*s*)^, the choice of power *p*, and the interval of relative age range will result in an actual equality:

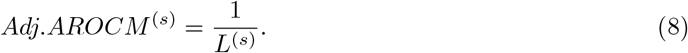

### Invariants

Our mathematical framework shows that equation (8) is a consequence from another major finding from our study: the Pearson correlation between methylation levels and age in selected chromatin states, Cor(*Methyl, Age*), does not exhibit a strong correlation with maximum lifespan. In particular, the age correlation in short-lived species will be similar to that in long-lived species. While we do not call the age correlation an “invariant,” our studies show that it does not relate to maximum lifespan across mammals (Figure 6).

### Development versus aging

Previous research in humans has indicated that the rate of change during development surpasses that during postnatal development [18, 27]. Our large mammalian data set corroborates this finding for a majority of mammalian species (Figure 4). Importantly, we demonstrate that the the rate of change in young is proportional to that in old animals (Figure 4). This latter discovery can be understood from two perspectives: mathematical and biological. In mathematical terms, it is a result (equation 26) of the continuous changes in cytosine methylation levels at particular locations, from development through old age. In fact, we offer explicit formulas that describe the monotonically increasing relationship between methylation and age across the life course (life course equations 27 and 28). From a biological standpoint, the connection between *AROCM*_*young*_ and *AROCM*_*old*_ reinforces deterministic theories of aging, suggesting a link between developmental changes and those occurring in later life [4, 14, 15, 21, 29]. Future versions of our life course equations might require modifications to accommodate for a potential rejuvenation phase early in development [30].

### Limitations and future research

Our research has multiple limitations, which stem from the inherent challenges in reliably calculating a rate of change. Careful examination of outliers in all strata is essential to reliable rate of change estimates, which we only ensured for the top chromatin states. To tackle these issues, we developed an outlier removal algorithm. Without this algorithm, the empirical data aligns far less with the mathematical formulas (Supplementary Figure S11). A notable limitation in our study is the wide fluctuation in age ranges across various strata (leading to different values of *SD*(*R*)). This can significantly influence the estimates of the rate of change. To address this, we introduced adjusted estimates for the rate of change. However, these adjustments are not without issues, including the risk of overfitting and challenges in interpretation. We recommend presenting both the unadjusted and adjusted rate of change estimates in findings, as done in our research. Our results remain consistent whether using adjusted or unadjusted estimates. Other limitations relate to potential confounders such as body size, tissue type, and phylogenetic relationships. Our multivariate model analysis partially addresses these concerns. Our results indicate that the inverse relationship between AROCM and mammalian lifespan persist even after adjusting for these confounders. A promising avenue for research is examining the disparities in methylation rate changes between genders. Additionally, it would be fascinating to investigate potential interventions influencing the AROCM. Many interventions influencing the lifespan of mice have been linked to alterations in age-associated methylation [24, 30, 35, 39, 46, 52]. While we present essential equations that connect epigenetic aging rates to maximum lifespan, this mathematical framework also holds significant relevance for numerous other aging biomarkers.

## Methods

### Methylation Platform

Both dog and mammalian data were generated using the same HorvathMammalMethylChip40 platform, which offers high coverage of approximately 36K conserved CpGs in mammals [2]. To minimize technical variation, all data were generated by a single lab (Horvath) using a single, consistent measurement platform. Preprocessing and normalization were performed using the SeSaMe method to define beta values for each probe [55]. The chip manifest file is available on the Gene Expression Omnibus (GEO) platform GPL28271 and on our Github page [2]).

### Chromatin states

Following the pan-mammalian aging study of the Mammalian Methylation Consortium, we grouped the CpGs into 54 universal chromatin states that were covered by at least 5 CpGs each [34]. These 54 chromatin states encompass those associated with both constitutive and cell-type-specific activity across a variety of human cell and tissue types [51]. In their 2022 study, Vu and Ernst employed a hidden Markov model approach to generate a universal chromatin state annotation of the human genome. This was based on data from over 100 cell and tissue types sourced from the Roadmap Epigenomics and ENCODE projects. These chromatin states are characterized in relation to 30 histone modifications, the histone variant (H2A.Z), and DNase I hypersensitivity measurements. We and others have previously found that strong age-related gain of methylation can be observed in bivalent promoter states and other states that are bound by Polycomb group repressive complex 2 (PRC2 binding sites) [34, 37, 41, 47]. To facilitate a detailed analysis of PRC2 binding, we split each chromatin state into 2 subsets denoted by “StateName+” and “StateName–” according to PRC2 binding (+ for yes and – for no). For example the “BivProm2+” is the set of 552 CpGs that reside in bivalent chromatin state 2 and are bound by PRC2 (Supplementary Table 5).

### Dog methylation data

We analyzed methylation profiles from *N* = 742 blood samples derived from 93 dog breeds (Canis lupus familiaris). Primary characteristics (sex, age, average life expectancy) for the breeds utilized are presented in Supplementary Table 1. Standard breed weight and lifespan were aggregated from several sources as detailed in [28]. We created consensus values based on the American Kennel Club and the Atlas of Dog Breeds of the World. Lifespan estimates were calculated as the average of the standard breed across sexes, compiled from numerous publications consisting primarily of surveys of multi-breed dog ages and cause of death from veterinary clinics and large-scale breed-specific surveys, which are often conducted by purebred dog associations. Sources for median lifespan per dog breed are reported in [28]. We calculated the maximum lifespan for dog breeds by multiplying the median lifespan with a factor 1.33, i.e. *MaxLifespan* = 1.33 ∗ *MedianLifespan*. Our results are qualitatively unchanged if other multipliers are used. Detailed values on the dog breeds are reported in Supplementary Table Median lifespans of the 93 breeds ranged from 6.3 y (Great Dane, average adult breed weight = 64 kg) to 14.6 y (Toy Poodle, average adult breed weight = 2.3 kg). Median lifespan estimates were based on the combined findings of multiple large-scale breed health publications, utilizing the median and maximum ages for each breed.

We identified 3 dog breeds (Otterhound, *n* = 4; Weimaraner *n* = 3; Saint Bernard Dog *n* = 2) as outlier strata for which the rate of change in methylation were not meaningfully estimated according to the following exclusion criteria. In addition to their small sample sizes, the age ranges are poor all with SD (**R**) *<* 0.1, resulting in extreme AROCM values *>* 0.5 (Supplementary Table 1). The remainder of dog breeds all had AROCM values no larger than 0.34. In summary, the following criteria should be considered for our dog data or other similar data in the future.

1. Small sample size, i.e. *n <* 3.
2. Low standard deviation in relative age, i.e. SD (**R**) *<* 0.1.
3. Bad linear regression fit of AROCM, i.e. *R*^2^ *<* 0.2.

### Selection of mammalian species/tissue strata

The raw data included 249 species-tissue strata from 133 unique species (Supplementary Figure S11). We selected strata with sufficient sample sizes and no influential outliers. Similar to the dog data, we excluded strata for the followings reasons.

1. Small sample size of *n <* 3.
2. Low standard deviation in relative age, i.e. SD (**R**) *<* 0.06, to avoid strata with constant ages.
3. Strata with AROCM values out of range (estimate *<* − 1 or *>* 10) were omitted if the values in derived/adjacent age intervals were not outliers. Toward this end, derived AROCMs were calculated for different age intervals within the same species/tissue stratum. For example, a severely outlying value *AROCM* [0, 0.1 ∗ *L*] was declared an outlier if both *AROCM* [0, 0.2 ∗ *L*] and *AROCM* [0, 0.3 ∗ *L*] fell within the range (−1, 10) for the same stratum.

To obtain additional strata for the dog data, we opted for a more lenient SD(R) cutoff. Given the greater number of strata in the mammalian data compared to the dog data (229 versus 94), we selected a less strict SD(R) threshold of 0.06 instead of 0.1 to ensure sufficient strata for analysis.

### Mammalian methylation data strata

Our data involved 133 unique mammalian species. For many species, several tissue types were available. The species characteristics such as maximum lifespan come from an updated version of the anAge data base [16, 23]. We analyzed *S* = 229 different species/tissue strata defined on the entire age range [0, *L*] (Supplementary Table 3). Out of the 229 strata, 100 involved blood, 73 skin, 26 brain, and 15 liver. Fewer strata were available for other age ranges. For example, *S* = 128 for the young age group (defined by [0, 0.1 ∗ *L*]) and *S* = 221 for the old age group (defined by [0.1 ∗ *L, L*]).

### Adjusted rate of change and adjusted correlation

The relative age *R*, defined as the ratio of age to maximum lifespan, is critical in disentangling the relationship between rates of change and maximum lifespan. In many real datasets, the standard deviation of relative age, SD (*R*), varies across species due to uneven sampling, which may primarily include young or old animals in some species. This variability in SD (*R*) often dilutes the signal between the rate of change and maximum lifespan, while affecting Cor(*Methyl, Age*), as evidenced in our simulation and empirical studies. Adjusting for SD (*R*) amplifies the inherent biological signal in both measures. Applying these formulas to the methylation data allowed us to present fundamental equations that link the rate of change in methylation in specific chromatin states (e.g. bivalent regulatory regions) to maximum lifespan in mammals.

Here we present a mathematical formalism that links three measurements: the rate of change in the biomarker across the life course, Pearson correlation between age and the biomarker, and the standard deviation of relative age. In most empirical data sets, the standard deviation of relative age is correlated with the Pearson correlation between age and the biomarker (Supplementary Figure S2), which reflects the idiosyncrasies of the sample collection. The standard deviation of age has a confounding effect on both the rate of change and the correlation between a biomarker and age. To study and to eliminate this confounding effect, we introduce two new concepts: *adjusted* rate of change and *adjusted* Pearson correlation. We present mathematical propositions describing the conditions under which strong relationships between rate of change and lifespan can be observed.

In the following, we derive general equations that link the rate of change (also known as gradient or slope) of any continuous biomarker of aging (denoted as *M* ∈ *R*) to the species maximum lifespan. For example, *M* could denote mean methylation in a particular chromatin state. Assume **M** = (*M*_1_, ‥*M*_*n*_) and **A** = (*A*_1_, ‥, *A*_*n*_) are two numeric vectors of *n* samples for the biomarker *M* and the Age variable. We will be using the following definitions surrounding the sample mean, sample variance, sample covariance, and Pearson correlation.

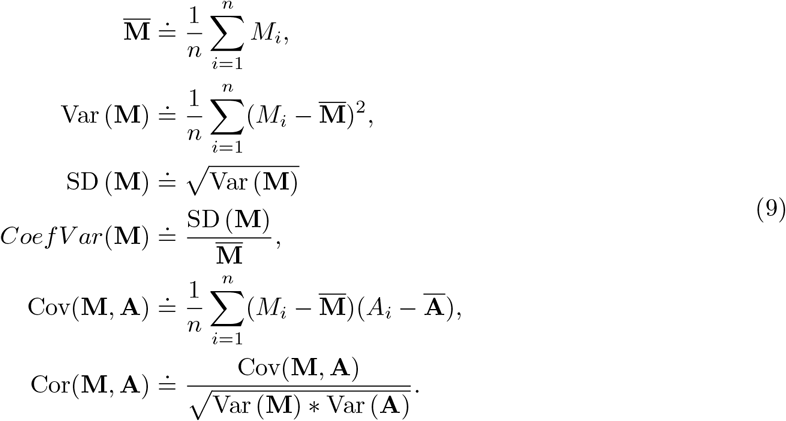

Next, we define the rate of change, ROC(**M**|**A**), as the change in *M* resulting from a 1-year increase in age (calendar age in units of years). Statistically speaking, the rate of change, ROC(**M**|**A**), is the slope/coefficient *β*_1_ in the univariate linear regression model below,

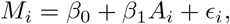

where the index *i* refers to the i-th tissue sample and the expected value of the error term *ϵ*_*i*_ is assumed to be zero. The rate of change can be estimated by the least squares or the maximum likelihood estimator, 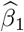. Furthermore, it can be expressed in terms of the Pearson correlation coefficient and standard deviations as follows

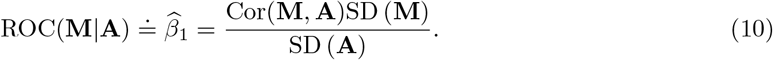

To arrive at a unit-less biomarker, which lends itself for comparisons with other biomarkers, we standardize **M** to have mean zero and standard deviation one, by scaling it as below,

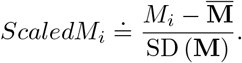

Using SD (**ScaledM**) = 1, equation (10) becomes

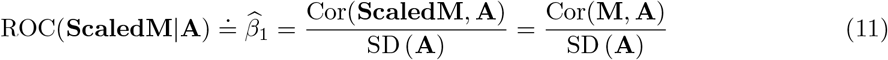

where the latter equation used the fact that the Pearson correlation, Cor, is invariant with respect to linear transformations. To reveal the dependence on species maximum lifespan, it is expedient to define relative age as the ratio of age and maximum lifespan:

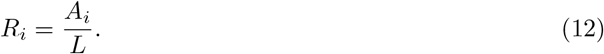

Since the standard deviation is the square root of the variance, one can easily show that SD (**A**) = SD (**R**) ∗ *L*. Combining equations (11) and (12) results in

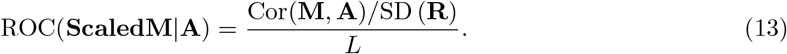

Since Pearson’s correlation is scale-invariant, the following equality holds and we will use them interchangeably, Cor(**M, A**) = Cor(**ScaledM, A**) = Cor(**M, R**).

**Proposition 1** (Relationship between ROC and Lifespan). *If the following condition holds across all strata*,

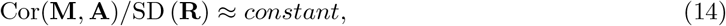

*then equation (13) implies*

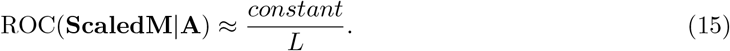

In empirical data, the strong condition (14) is not always satisfied, see e.g. Figure S2d,e. This is because of the reasons mentioned previously due to sampling bias, sample sizes, or imperfect age distributions. But we propose a simple adjustment to formulate a weaker condition that is more realistic and adaptive to real data, which leads to a similar conclusion as equation (15). All we need is to rewrite equation (13) as below

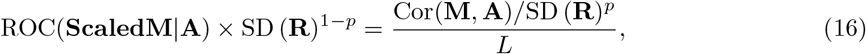

which multiplies both sides by SD (**R**)^1−*p*^ with a power parameter *p*. Naturally, we define:

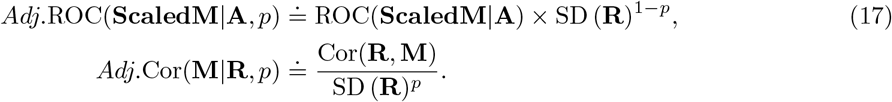

Note that if SD (**R**) remains constant across strata, indicative of a perfect design, the adjustment essentially involves multiplying or dividing by a constant, irrespective of the power *p*. This means the adjustment leaves the relationship between ROC and lifespan unchanged. On the other hand, if SD (**R**) fluctuates across strata—indicative of an imperfect study design—the adjustments have the potential to enhance the signal. Further note that Adj.ROC becomes the standard definition of the ROC with *p* = 1. On the other hand, *p* = 0 implies that *Adj*.Cor(**M R**, *p*) = Cor(**M, R**). The novel terminology has several advantages. To begin with, equation (16) can be succinctly written as follows

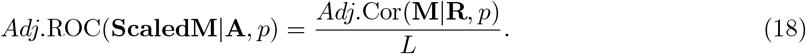

The goal is to derive conditions when equation (18) satisfies the following

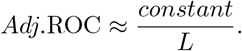

The approximation “≈ “means that there is a high linear correlation across different strata on the log scale. We start with the log transformed version of equation (18):

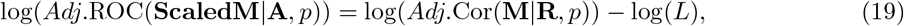

where we assume that the natural logarithm (log) is applicable, i.e. the adjusted ROC and the adjusted correlation take on positive values.

The above mentioned definitions and equations apply to each stratum (e.g. dog breeds). Assuming there are *S* total strata, we introduce a superscript in various quantities e.g. we write *L*^(*s*)^, and *Adj*.Cor(**M**^(*s*)^ | **R**^(*s*)^, *p*), where *s* = 1, 2, ‥, *S*. Define the following 3 vectors that have *S* components each

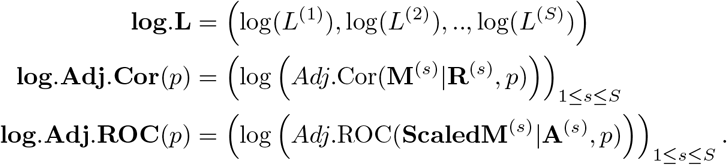

For each vector on the left hand side, we can form the sample mean and sample variances across *S* strata,

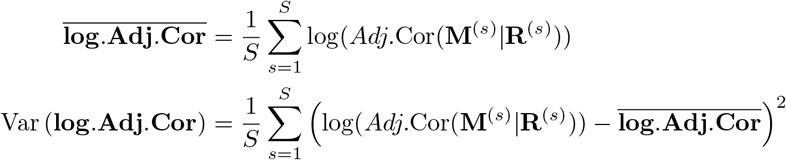

The key condition is the following,

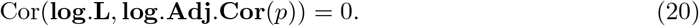

**Remark 1** The condition Cor(**log.L, log.Adj.Cor**(**p**)) = 0 holds when species lifespan and the *Adj*.Cor(*p*) are independent across strata. Our methylation data suggest that this condition is approximately satisfied for certain chromatin states.

We are now ready to state the main proposition.

**Proposition 2** (Linear relationship between log.Adj.ROC and log.L). *If (C1) holds and the coefficient of variation in Adj*.Cor(*p*) *is much smaller than the coefficient of variation in L, i*.*e*.

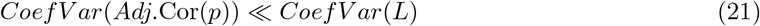

*then*

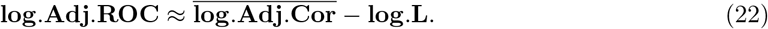

The proof of this proposition and related properties can be found in **Supplementary Methods** separately. Exponentiating both sides of equation (22), we arrive at

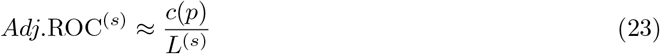

where 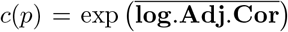 is some constant. The choice of the parameter *p* will be discussed below. While real data may not consistently satisfy Condition C1, equation (23) frequently remains roughly accurate. This pattern suggests that equation (23) might be derived under more lenient assumptions.

### Criteria for choosing the power *p* in the adjustment

Our above mentioned equations make use of the parameter *p* that underlies our definitions of the adjusted correlation and the adjusted ROC. A choice of *p* = 1 results in standard (non-adjusted) versions of the ROC but it can be advantageous to choose a lower value of *p* for the following three reasons.

First, Proposition 2 states that a strong linear relationship between **log.Adj.ROC** and **log.L** holds if one chose *p* such that it minimizes the coefficient of variation function *C*(*p*) = *Coef V ar* (*Adj*.Cor(*p*)). Since the coefficient of variation is more sensitive to outliers, we also use a robust version known as the quartile coefficient of dispersion (QCOD):

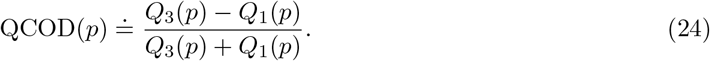

where *Q*_1_(*p*) and *Q*_3_(*p*) denote the first and third quartile of the distribution of *Adj*.Cor(*p*). In our empirical studies, we chose *p* so that it minimized QCOD (equation 24), i.e.

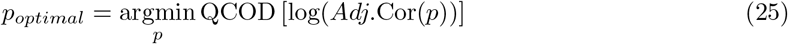

Using the QCOD-based criterion, we determined *p*_*optimal*_ = 0.1 for our dog data and *p*_*optimal*_ = 0.25 for our mammalian methylation dataset (refer to Supplementary Figure S1). Had we employed the coefficient of variation in place of the QCOD, our choice of *p* would have been consistent across both datasets, as depicted in Supplementary Figure S1. This alignment between the coefficient of variation and QCOD is well-documented in statistical literature, as cited in [6, 38].

The second reason for choosing the power *p* relates to an undesirable correlation between the age correlation Cor(**M, A**) and the standard deviation of relative age, SD (**R**) (Figure S2). Our simulation studies suggests that this positive correlation results from an imperfect sample ascertainment/study design. This can be mitigated against by choosing *p* so that the *adjusted* age correlation *Adj*.Cor(*M*| **R**, *p*) exhibits a weaker correlation with SD (**R**). In the mammalian data, *p* = 0.25 leads to non significant correlation between *Adj*.Cor(*M* | **R**, *p*) and SD (**R**) (Figure S2g,h).

Third, our simulation studies that aimed to emulate our mammalian lifespan data indicate that with large sample sizes per stratum *Adj*.Cor(*M* | **R**, *p*) converges to 1 for *p* = 0.25 (Supplementary Methods). Overall, these results suggest that *p* = 0.25 is a good choice for our mammalian methylation study.

### Relation between *AROCM*_*young*_ and *AROCM*_*old*_

Here, we provide an outline on how to derive a relationship between the rate of change in young animals and that in older ones in the s-th species tissue stratum, i.e.

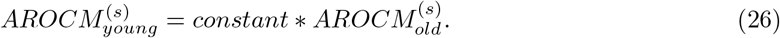

where *constant* denotes a number. We start out with commenting on our definition of relative age. When dealing with prenatal samples (whose chronological ages take negative values), it can be advantageous to slightly modify the definition of relative age as 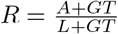, by including gestation time (GT) to avoid negative relative ages. For simplicity, we will assume that our data only contains postnatal samples, allowing us to define relative age as 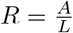. Empirically, we find that non-linear relationship between *ScaledM* and relative age in each stratum can be approximated using the following function:

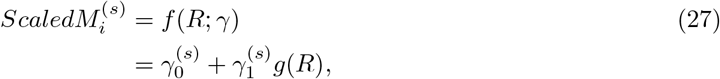

where 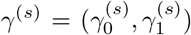are stratum-specific constants. Our empirical studies demonstrate that the following log linear function fits the data quite well.

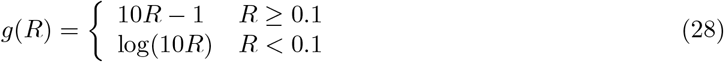

Note that the first derivative of *g*() is given by

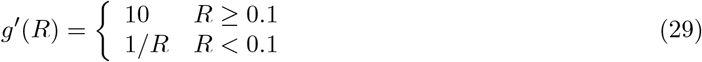

Assuming a linear relationship between *ScaledM* and *A* and a suitably chosen point *A*_0_, one can approximate AROCM as follows

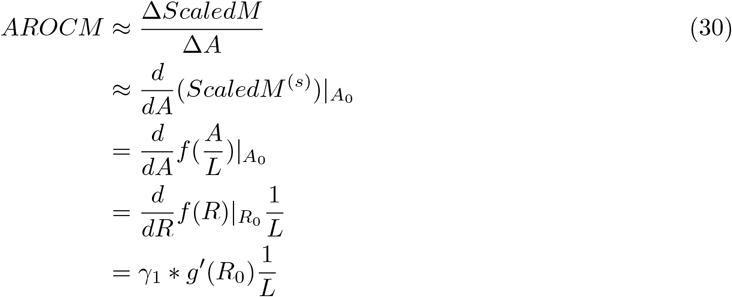

where *R*_0_ = *A*_0_*/L* and we used the chain rule of calculus. We define the AROCM in young and old animals as the first derivative evaluate at *A*_*young*_ and *A*_*old*_, respectively. These ages should be chosen so that the corresponding relative ages *R*_*young*_ and *R*_*old*_ take on values *<* 0.1 and *>* 0.1, respectively. With equations 29 and 30, we find

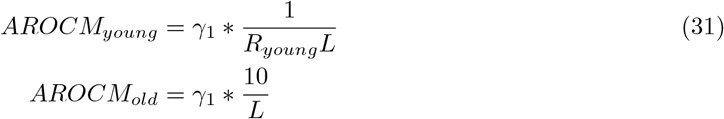

With superscripts denotidng the s-th species tissue stratum, the latter implies the following linear relationship between the two aging rates

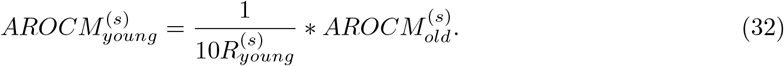

If 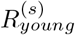 is roughly constant then the latter implies that 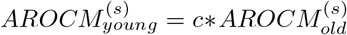. Empirically, we can verify the latter relationship (Figure 4). Across species-tissue strata, we find that *c* has a mean value 7.33 and standard deviation 6.8.

## Supporting information

Supplementary Table 1 Dog slopes

Supplementary Table 2 Chromatin States

Supplementary Table 3 AROCM Strata

Supplementary Table 4 Slopes By Species

Supplementary Table 5 CG list BivProm2+

## Data Availability

Individual level data from the Mammalian Methylation Consortium will be distributed via Gene Expression Omnibus once the primary citation of the Mammalian Methylation Consortium has been published [23]. The mammalian array platform is distributed by the non-profit Epigenetic Clock Development Foundation (https://clockfoundation.org/). The manifest file of the mammalian array and genome annotations of the CpGs can be found on Github (doi: 10.5281/zenodo.7574747).

## Software Availability

Our software is being distributed through GitHub, at github.com/feizhe/FundamentalEquations.

## Supplementary Figures

**Figure S1:**
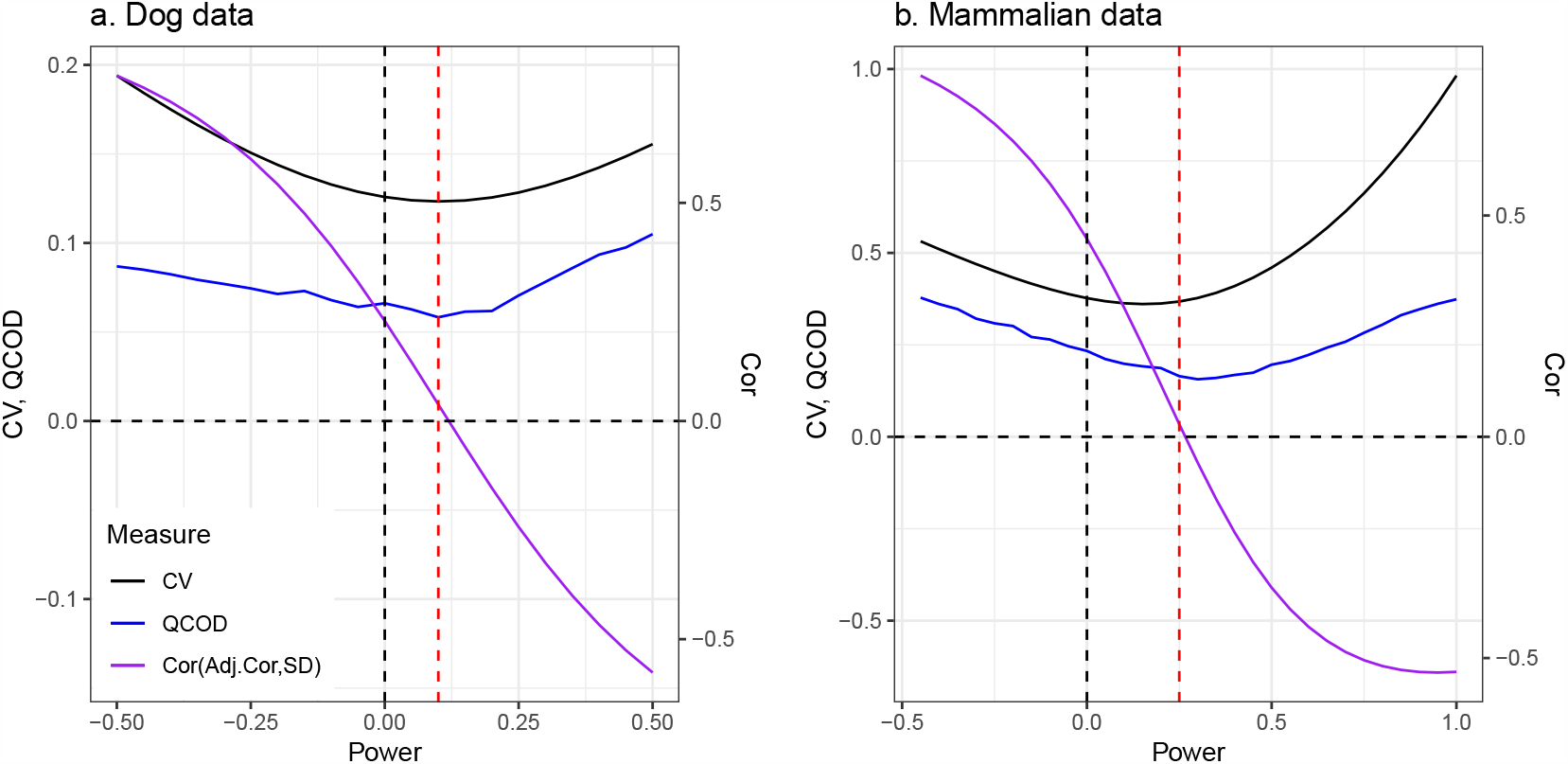
Optimal power for Adj.ROC in (a) Dog data and (b) Mammalian data. CV (Coefficient of Variation equation 9), QCOD (equation 24), and Cor(*Adj*.Cor, SD (*R*)) are plotted versus different powers. The optimal power (red line) is chosen to be 0.1 for the dog data and 0.25 for the mammalian data.

**Figure S2:**
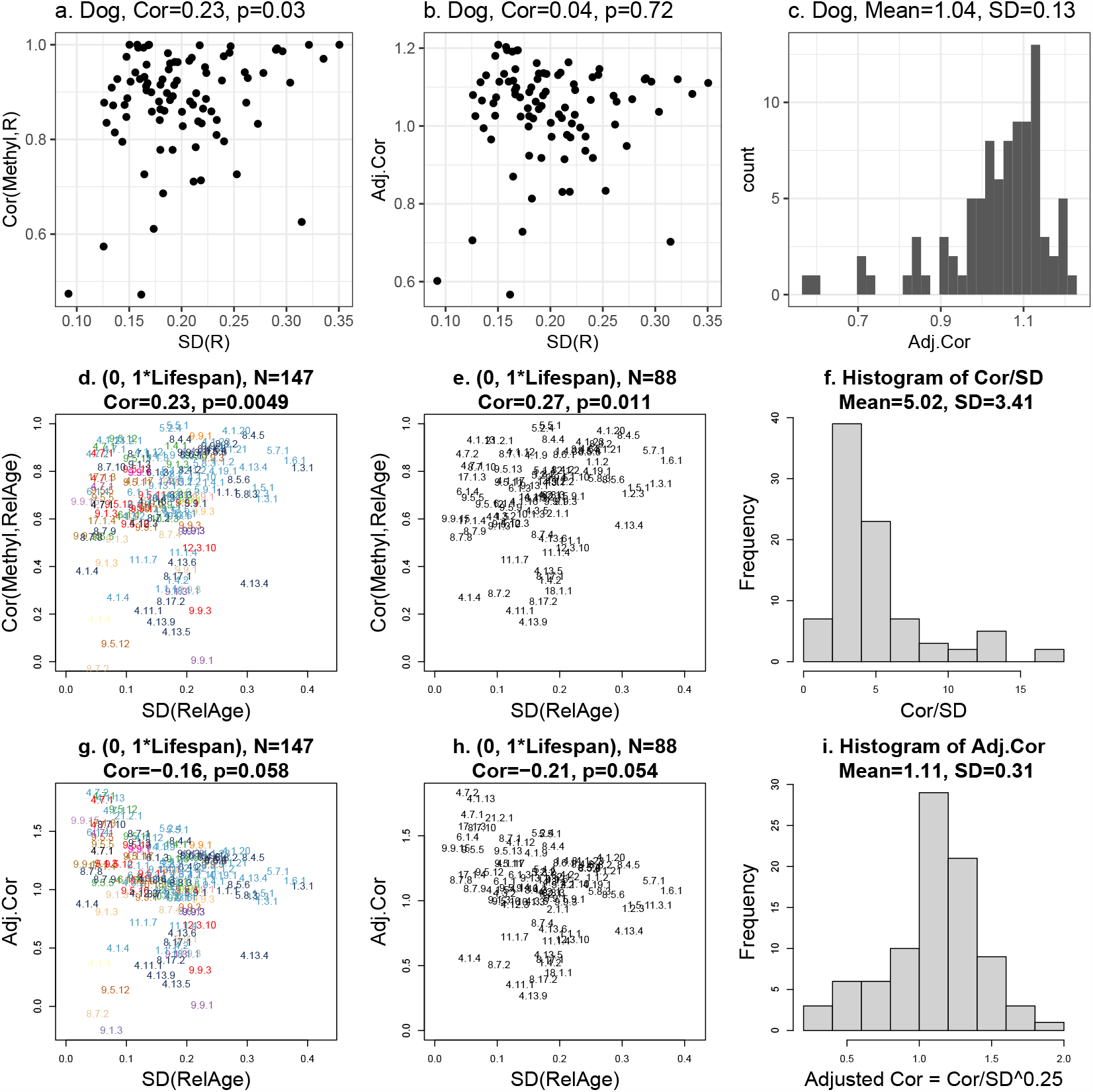
Age Correlation versus Standard Deviation of Relative Age. **(a-c)** dog data; **(d-i)** mammalian data. **(a**,**d**,**e)** age correlation Cor(*M, A*) and **(b**,**g**,**h)** the adjusted age correlation *Adj*.Cor against the standard deviation of relative age SD (**R**). Histograms of **(c)** Adj.Cor in dogs, **(f)** Cor*/*SD in mammals, and **(i)** Adj.Cor in mammals are also provided. Titles of scatter plots report Pearson correlation coefficients and associated Student T test p-values. Insignificant Pearson correlation test p values suggest adjusted age correlations do not correlate with SD (*R*), indicating adjusted correlation coefficients provide better resilience against sample ascertainment biases.

**Figure S3:**
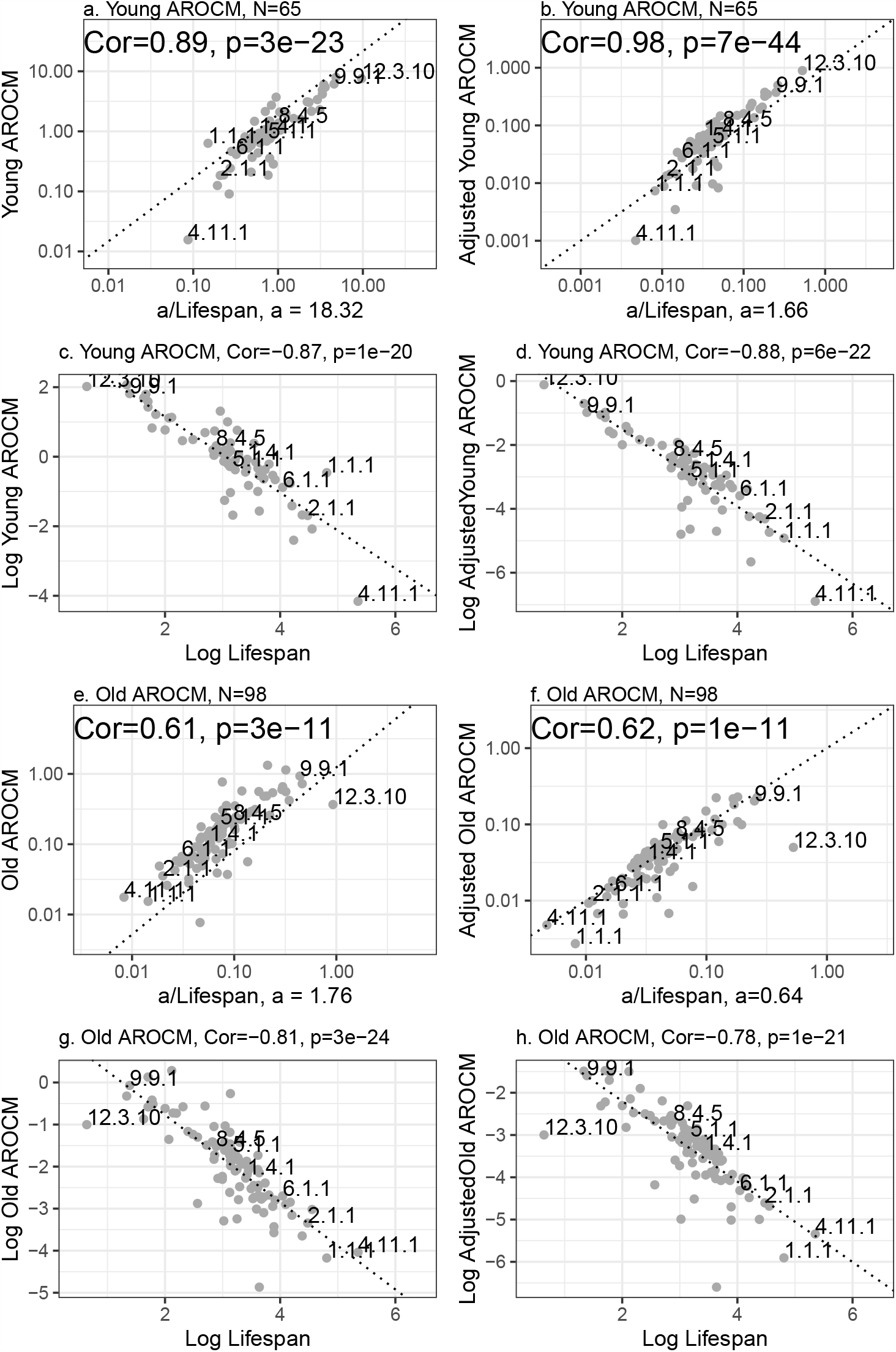
Young and Old AROCM versus max lifespan. This figure is analogous to Figure 5 but with Young and Old AROCMs. **a-d**. the sample size for Young AROCMs is *N* = 65; **e-h**. the sample size for Old AROCMs is *N* = 98.

**Figure S4:**
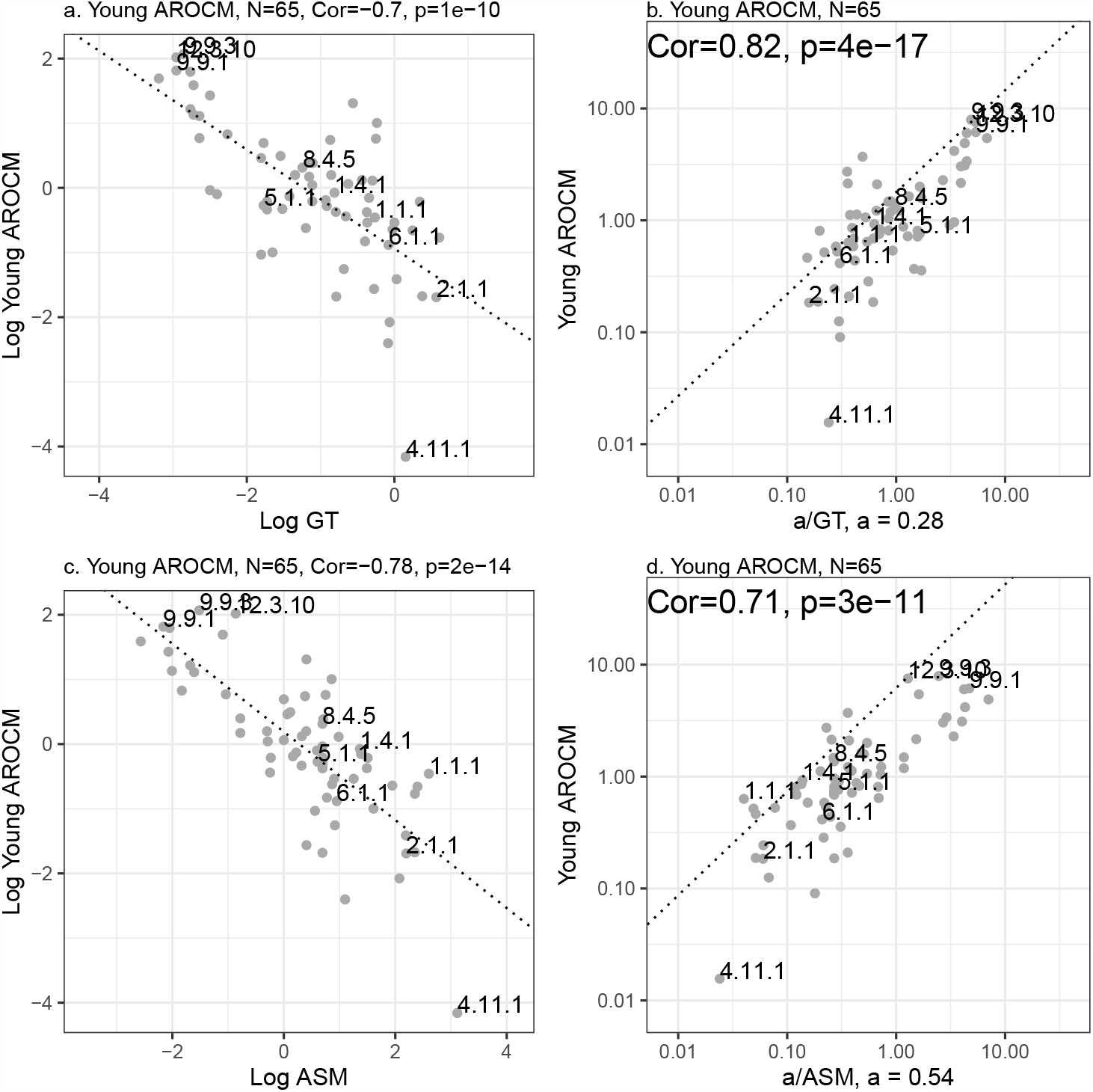
AROCM in young animals versus gestation time. Young AROCM[0, 0.1*L*] vs. Gestational Time (GT, panels a,b) and Age at Sexual Maturity (ASM, panels c,d).

**Figure S5:**
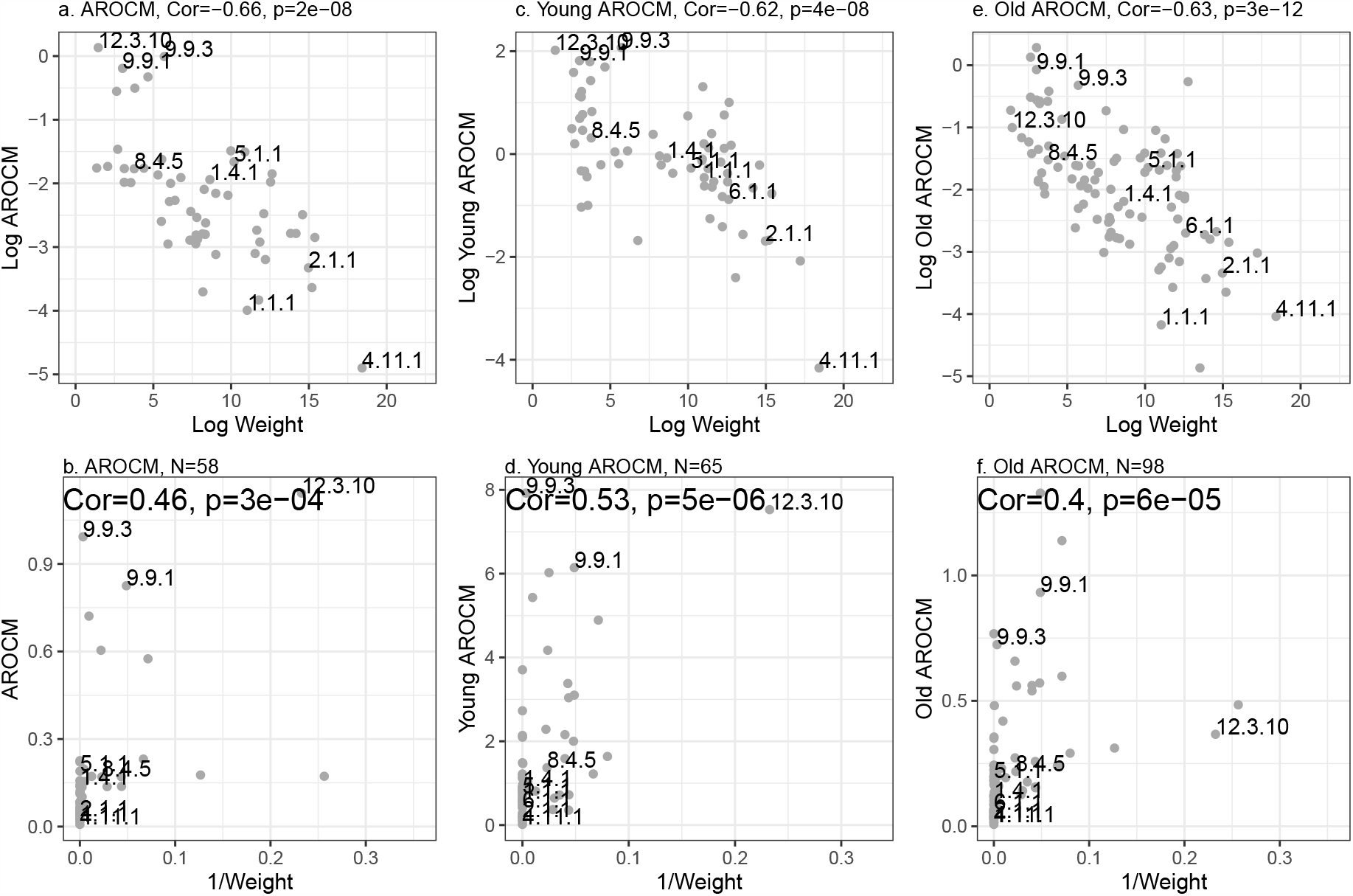
AROCM versus average adult weight in different age groups. Columns correspond to different age ranges: (a,b) all ages AROCM, (c,d) Young AROCM[0, 0.1*L*], and (e,f) old AROCM. The first row panels (b,c,e) present log transformed AROCM (y-axis) against average adult weight (x-axis). The second row panels (b,d,f) contrast untransformed AROCM values (y-axis) with 1*/Weight* (x-axis)

**Figure S6:**
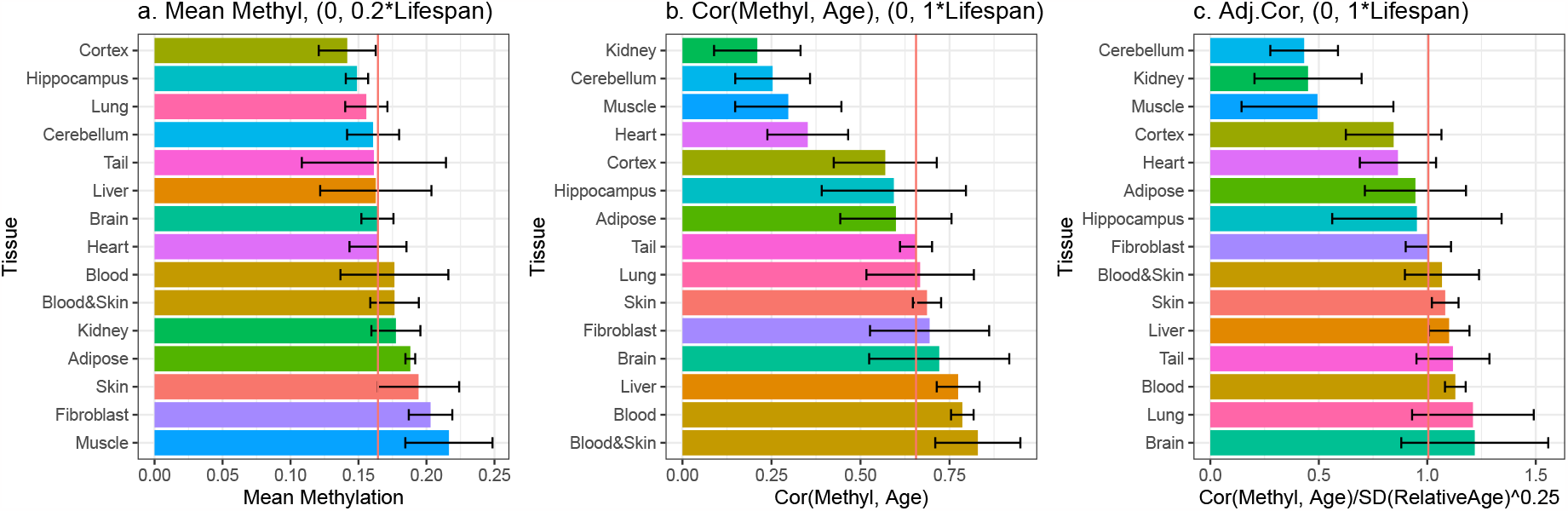
Age correlations in different mammalian tissue types. This figure depicts Mean Methylation, Cor(Methyl, Age), and Adjusted Cor in BivProm2+ across tissue types. Error bars signify 1 standard error from the median per tissue type, with the red vertical line indicating the overall median. Results, however, are confounded by species, prompting the use of multivariate regression models to explore tissue type effects (Table 1). The number of species within each tissue can be found in Supplement Table 3.

**Figure S7:**
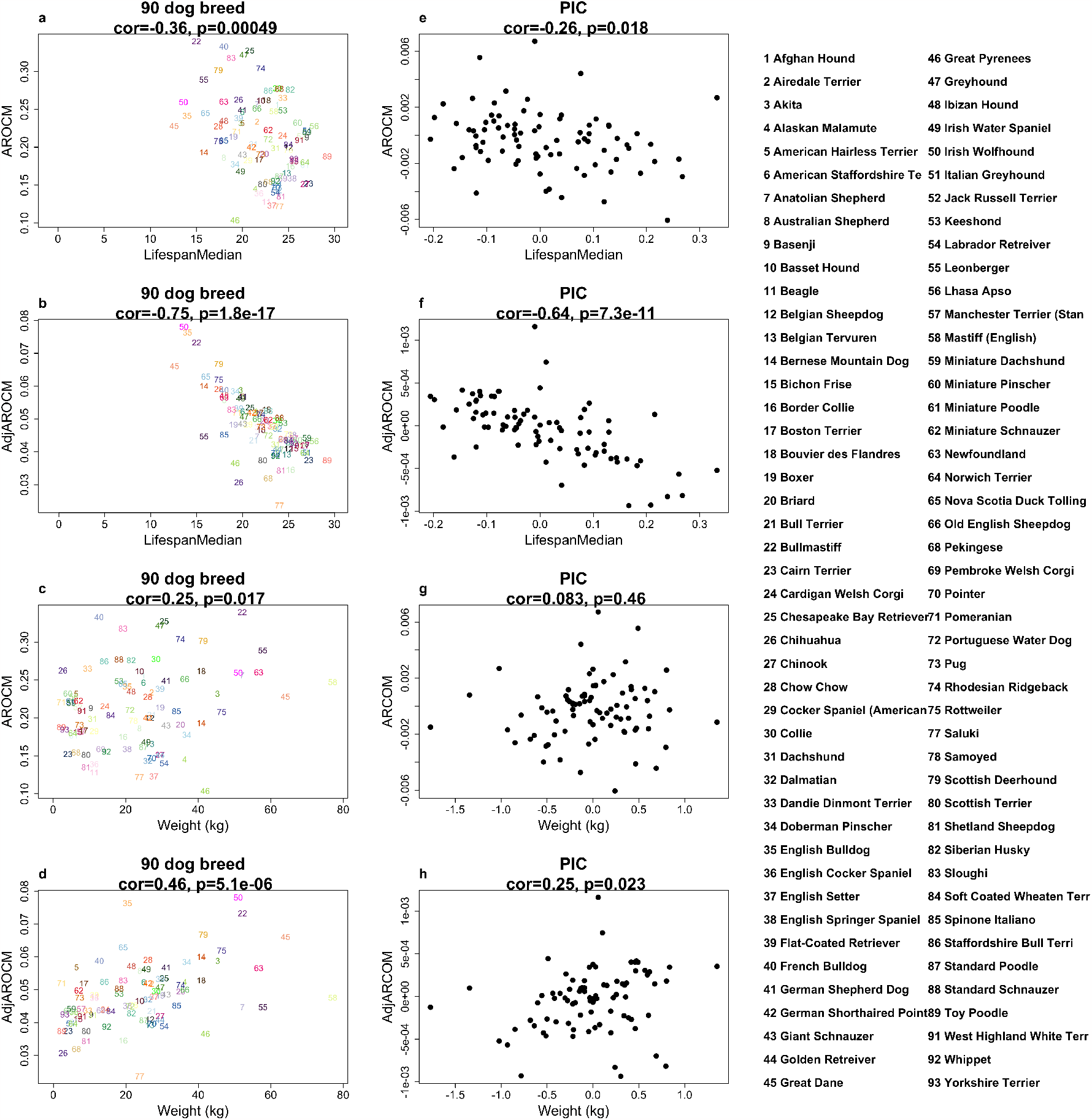
Phylogenetic independent contrasts of AROCM in dog breeds. Dog breed phylogenetic relationships were established as per [28]. The left panels (a,b,c,d) illustrate original variable relationships, while the right panels (e,f,g,h) present related results for phylogenetic independent contrasts (PICs, [19]). Accounting for these phylogenetic relationships via phylogenetic regression, both AROCM and Adj.AROCM retain significant lifespan correlations (e,f).

**Figure S8:**
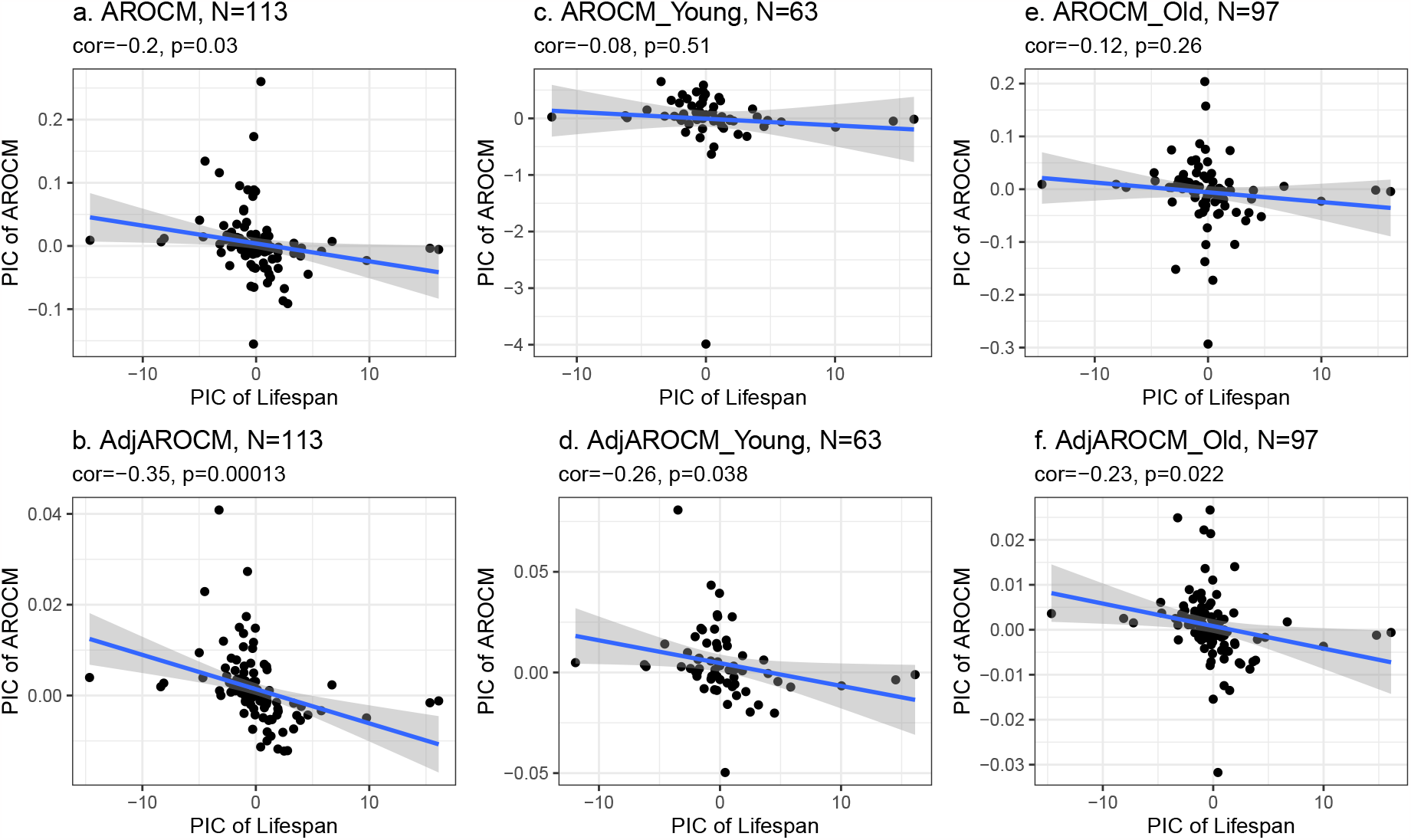
Phylogenetic independent contrasts of AROCM in mammalian species. The columns represent various age ranges: a,b) AROCM for all ages, c,d) young AROCM[0, 0.1*L*], and e,f) old AROCM. The first and second row panels (a,c,e; b,d,f) show results for unadjusted and adjusted AROCM, respectively. Despite considering the phylogenetic links between mammalian species, the three adjusted AROCMs maintain significant lifespan associations. The phylogenetic relationships between species come from TimeTree [32].

**Figure S9:**
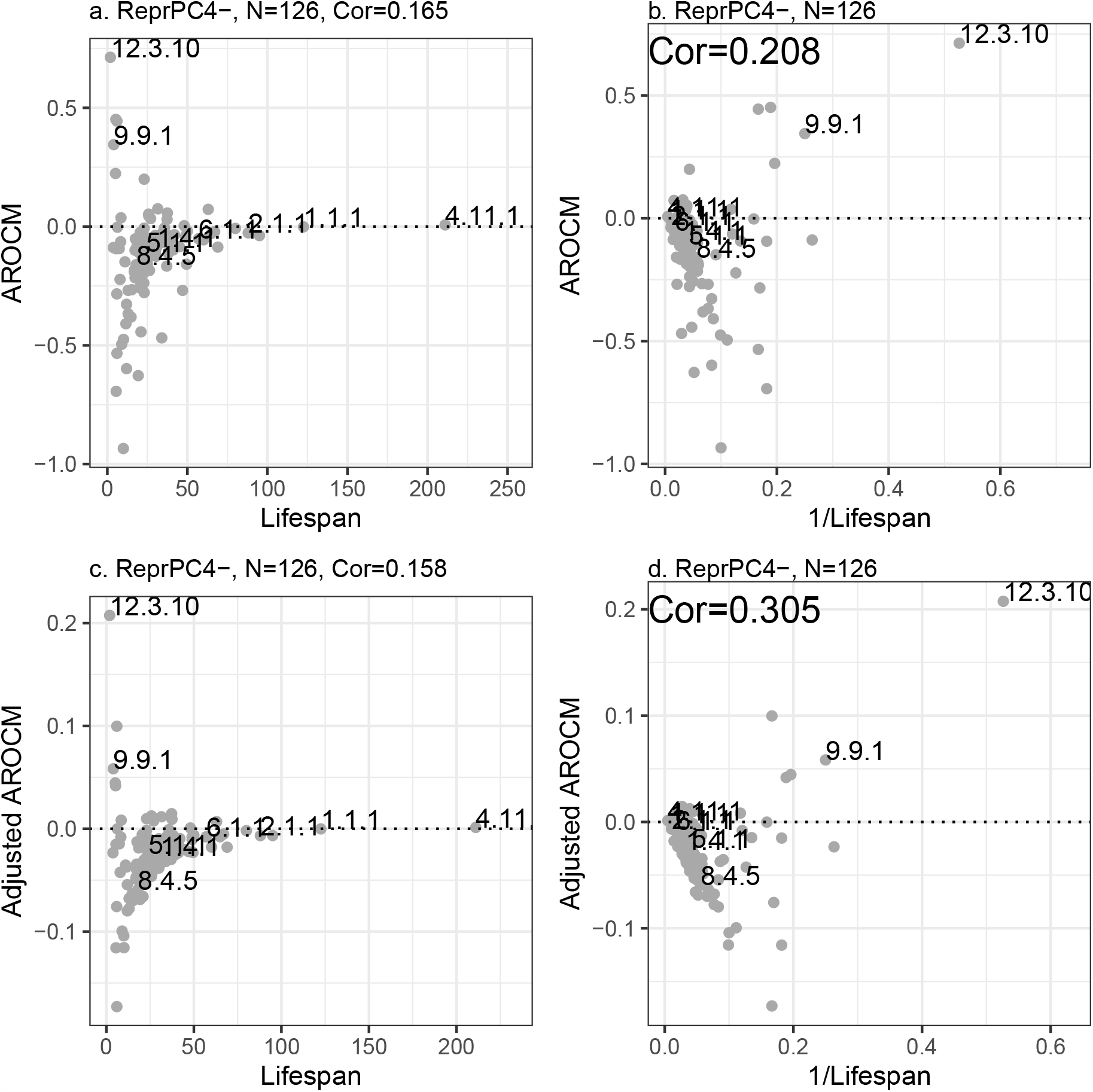
Non-significant association between AROCM and Lifespan in some chromatin states. Optimal power chosen to be 0.85 for ReprPC4-.

**Figure S10:**
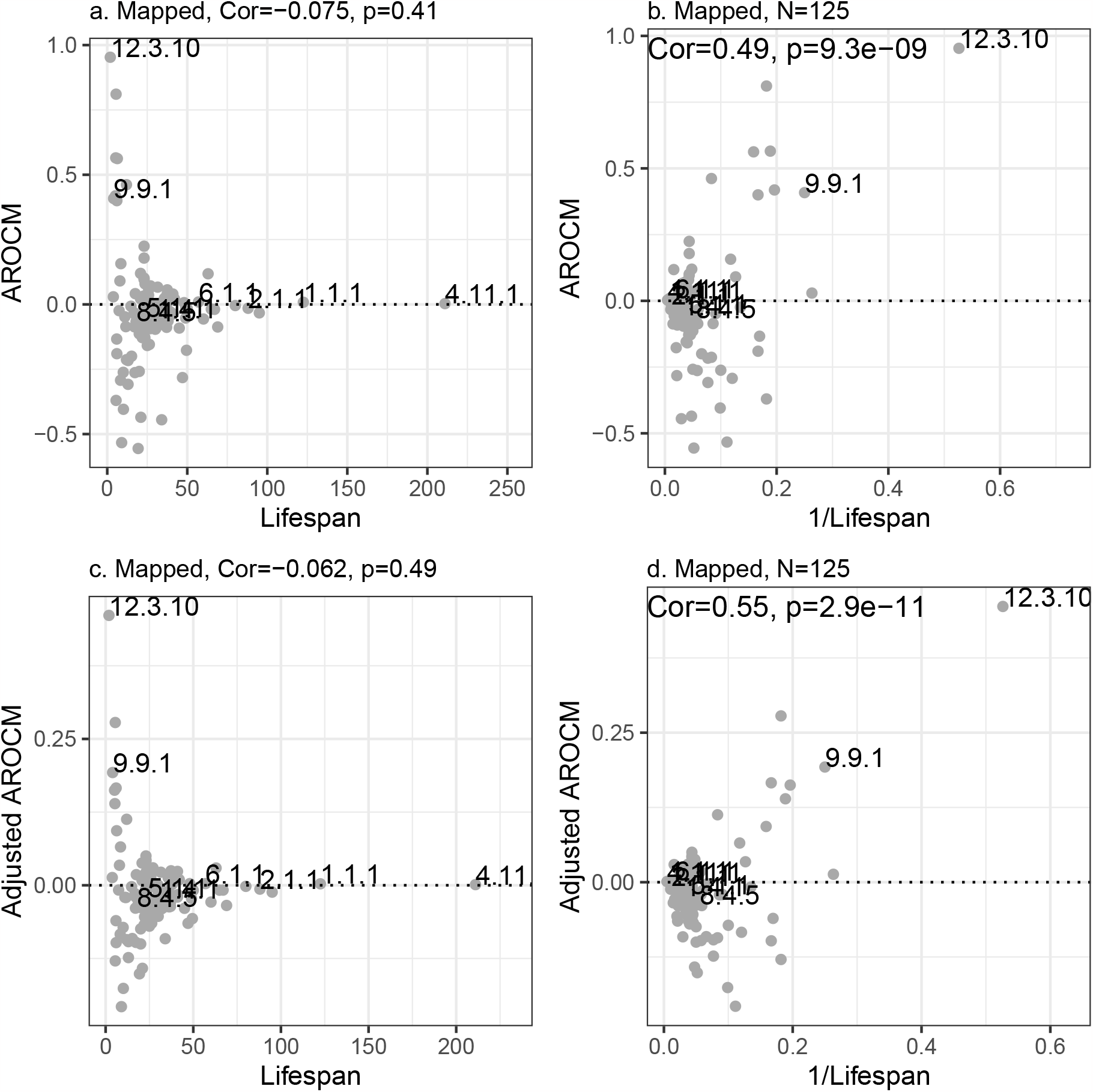
Non-significant association between AROCM and Lifespan in all CpG sites mapped to Eutherians and Marsupials. (*p* = 8970). Sample size *N* = 125. Optimal power chosen to be 0.5 for Adj.AROCM.

**Figure S11:**
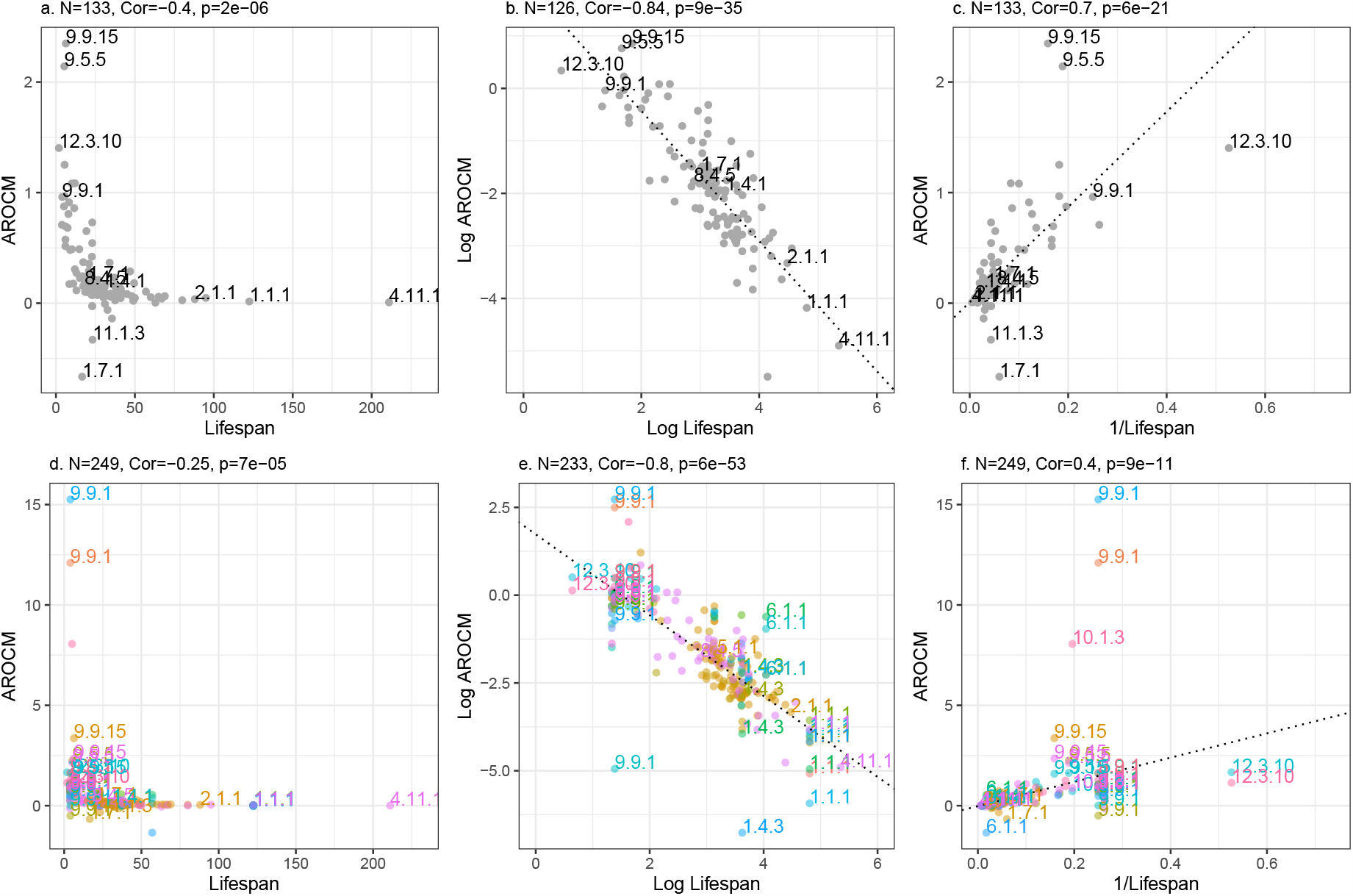
AROCM versus Lifespan with outliers in mammalian data. This figure is analogous to Figure 5 but with outliers not filtered by the criteria in Methods.

