## Supplementary Methods for "Fundamental equations linking methylation dynamics to maximum lifespan in mammals"

#### Simulation of AROCM

The fact that the number of animals per species and tissue stratum varies greatly affects both the correlation coefficients and standard deviations of different measures of interest within each stratum, as we will show using simulation studies.

Here we simulate data sets comprised of different species and tissue strata with varying sample sizes per stratum. Specifically, we use simulation studies to demonstrate that one can observe the following results:

- For stratum $s$ ($s = 1, ..., S$), define the Pearson correlation $Cor(ScaledMeth^{(s)}, \mathbf{R}^{(s)})$ and the standard deviation of relative age $\text{SD}(\mathbf{R}^{(s)})$. We find that these two quantities are correlated due to the different number of animals per stratum $n_s$.

- Adjusted Correlation, the ratio with a suitable power (e.g. $p = 0.25$),

$$Adj.Cor(p) = Cor(ScaledM^{(s)}, \mathbf{R}^{(s)})/\text{SD}(\mathbf{R}^{(s)})^p,$$

  is less dependent on the standard deviation $\text{SD}(\mathbf{R}^{(s)})$ than the correlation $Cor(ScaledM^{(s)}, \mathbf{R}^{(s)})$.

To demonstrate these results, we simulated different scenarios of data sets with $S = 200$, to mimic our real data. Each simulated stratum represents a different species whose maximum lifespan was randomly chosen to lie between 2 years (e.g. a shrew) and 250 years (e.g. long lived whales). We assumed that on a log scale, the species lifespan followed a uniform distribution, i.e., $\log L^{(s)} \sim Unif(\log 2, \log 250)$. The number of animals per stratum, $n_s$, was randomly sampled from a Poisson distribution whose mean value was set to $\mu$. We explored 3 different values of $\mu = 20, 100, 1000$. In other words, $\mu = 1000$ would result in an ideal data set comprised of 1000 animals per species on average. The chronological age of each animal was randomly chosen from a uniform distribution between 0 and maximum lifespan, $A_i^{(s)} \sim Unif(0, L^{(s)})$. For each animal, the relative age was defined as the ratio between age and maximum lifespan. The mean methylation level (Methyl) was simulated to be a linear function of Relative Age, $M_i^{(s)} = R_i^{(s)} + e_i^{(s)}$, where $e_i^{(s)} \sim N(0, 0.5^2)$. For samples within each stratum, we scaled the mean methylation levels so that it would have a mean of zero and a variance of 1, i.e. $ScaledM_i^{(s)} = \frac{M_i^{(s)} - Mean(M_i^{(s)})}{\text{SD}(M_i^{(s)})}$. Note that scaling does not change the correlation between mean methylation and age.

Within each stratum, we calculated the ratio $Adj.Cor(p) = Cor(ScaledM, \mathbf{R})/\text{SD}(\mathbf{R})^p$, where $p$ is the tuning parameter power. Next, we studied the mean value, standard deviation, and the distribution of this ratio across all strata, as well as its relationship with $\text{SD}(\mathbf{R}^{(s)})$ (Figure S12). Interestingly, the correlation between $\text{Cor}(ScaledMethyl, \mathbf{R})$ (y-axis) and $\text{SD}(\mathbf{R})$ (x-axis) remains around 0.3, independent of the average sample size per stratum (Figure S12 a-c), while the variances of these quantities strongly depend on the average sample size. However, the dependency between the Adjusted Correlation and $\text{SD}(\mathbf{R})$ is much weaker and not significant (Figure S12 d-f). The mean of Adjusted Correlation remains the same when the average sample size increases (Figure S12 g-i).

Next, we investigate the power term used to define the Adjusted Correlation,

$$\begin{aligned} Adj.AROCM =& AROCM * \text{SD}(\mathbf{R})^{1-p} \\ =& \frac{\text{Cor}(ScaledM, \mathbf{R})}{\text{SD}(\mathbf{R})^p} \frac{1}{Lifespan}. \end{aligned} \tag{1}$$

The goal is to optimize the ratio $Adj.\text{Cor}(p) = \text{Cor}(ScaledM, \mathbf{R})/\text{SD}(\mathbf{R})^p$ that whose Coefficient of Variation is minimized for certain power value, $CV(p) = \text{SD}(Ratio(p))/Mean(Ratio(p))$.

We simulated different scenarios with $S = 50, 100, 200, 1000$, and 2000, and checked how do the CVs change with respect to the power $p$ in $Adj.$Cor (Supplementary Figure S13). The optimal power is achieved at 0.25 where the Coefficient of Variation is minimized.

For each scenario, we simulated 1000 replicate data ($nsim = 1000$) and averaged the results. In the simulation scenario that mimics our real data, we find that empirically the mean of the ratio approximates 1 when $p = 0.25$. Here we simulate the strata sample sizes following $\log n \sim N(\mu_0, \sigma_0^2)$, where $\mu_0, \sigma_0$ are the empirical values from the real data.

### Additional Propositions

Recall the relationship

$$\log(Adj.\text{ROC}(\mathbf{ScaledM}|\mathbf{A}, p)) = \log(Adj.Cor(\mathbf{M}|\mathbf{R}, p)) - \log(L), \tag{2}$$

and the assumption

$$CoefVar(Adj.\text{Cor}(p)) \ll CoefVar(L) \tag{3}$$

**Proposition S1** (Correlation between log.Adj.ROC and log.L)**.** *If (C1) holds, then*

$$\text{Cor}\left(\mathbf{log.L}, \mathbf{log.Adj.ROC(p)}\right) = \frac{-1}{\sqrt{1 + \text{Var}\left(\mathbf{log.Adj.Cor}\right)/\text{Var}\left(\mathbf{log.L}\right)}}$$

**Remark 2** This proposition implies that Lifespan and Adj.ROC follow a nearly perfect inverse linear correlation on the log scale (Cor $\approx -1$) if $\text{Var}(\mathbf{log.Adj.Cor}) \ll \text{Var}(\mathbf{log.L})$. The latter condition is typically satisfied in real data as the range of lifespans across strata is often much larger than the Adj.Cor values, which is the case for our data from the mammalian methylation consortium.

*Proof.* Denote vectors $\mathbf{x} = \mathbf{log.L}$ and $\mathbf{y} = \mathbf{log.Adj.ROC}$. With equation (2) we find that the covariance

$$\text{Cov}(\mathbf{x}, \mathbf{y}) = \text{Cov}(\mathbf{log.L}, \mathbf{log.Adj.Cor}) - \text{Cov}(\mathbf{log.L}, \mathbf{log.L})$$

By assumption the first term is zero, which entails that

$$\text{Cov}(\mathbf{x}, \mathbf{y}) = -\text{Cov}\left(\mathbf{log.L}, \mathbf{log.L}\right) = -\text{Var}\left(\mathbf{x}\right)$$

Similarly, $\text{Cov}(\mathbf{log.L}, \mathbf{log.Adj.Cor}) = 0$ implies that

$$\begin{aligned}\text{Var}\left(\mathbf{y}\right) &= \text{Var}\left(\mathbf{log.L}\right) - 2 * \text{Cov}(\mathbf{log.L}, \mathbf{log.Adj.Cor}) + \text{Var}\left(\mathbf{log.Adj.Cor}\right)\\ &= \text{Var}\left(\mathbf{log.L}\right) + \text{Var}\left(\mathbf{log.Adj.Cor}\right)\end{aligned}$$

Thus, the assumption implies that

$$\begin{aligned}\text{Cor}(\mathbf{x}, \mathbf{y}) &= \frac{\text{Cov}(x, y)}{\sqrt{\text{Var}\left(\mathbf{x}\right)\text{Var}\left(\mathbf{y}\right)}}\\ &= -\frac{\text{Var}\left(\mathbf{x}\right)}{\sqrt{\text{Var}\left(\mathbf{x}\right)\left(\text{Var}\left(\mathbf{x}\right) + \text{Var}\left(\mathbf{log.Adj.Cor}\right)\right)}}\\ &= -\frac{1}{\sqrt{1 + \text{Var}\left(\mathbf{log.Adj.Cor}\right)/\text{Var}\left(\mathbf{log.L}\right)}}.\end{aligned}$$

□

The following proposition is a direct consequence of Proposition S1.

**Proposition S2** (Linear relationship between log.Adj.ROC and log.L)**.** *If (C1) holds and the ratio*

$$\text{Ratio}(p) = \frac{\text{Var}\left(\mathbf{log.Adj.Cor}(p)\right)}{\text{Var}\left(\mathbf{log.L}\right)} \approx 0, \tag{4}$$

*then*

$$\mathbf{log.Adj.ROC} \approx \overline{\mathbf{log.Adj.Cor}} - \mathbf{log.L}. \tag{5}$$

*Proof.* By (2), the LHS and RHS of (5) have the same mean, i.e. $\overline{\mathbf{log.Adj.ROC}} = \overline{\mathbf{log.Adj.Cor}} - \overline{\mathbf{log.L}}$. Proposition S1, combined with the assumption that $\text{Ratio}(p) \approx 0$, leads to the conclusion that $\text{Cor}(\mathbf{log.Adj.ROC}, \mathbf{log.L}) \approx -1$. Given that a Pearson correlation nearing negative one indicates an almost perfect linear relationship, this finalizes the proof. □

Proposition 2 is a variant of Proposition S2 that can be more easily verified in most real data sets. Its proof is stated below.

*Proof.* In the following, we will show that the assumption (equation 3) implies equation (4) in Proposition S2. We will use the following Delta method approximation for computing the variance of $f(X)$ of a random variable $X$,

$$\text{Var}\,(f(X)) \approx f'(\text{E}(X))^2 \text{Var}\,(X),$$

where $\text{Var}\,(X)$ and $\text{E}(X)$ denote the variance and expectation of $X$, respectively. With $f(x) = \log(x)$, $f'(x) = 1/x$ and $X = \mathit{Adj}.\text{Cor}(p)$, the above approximation results in

$$\text{Var}\,(\log(\mathit{Adj}.\text{Cor}(p))) \approx \frac{\text{Var}\,(\mathit{Adj}.\text{Cor}(p))}{\text{E}(\mathit{Adj}.\text{Cor}(p))^2} = CoefVar^2(\mathit{Adj}.\text{Cor}(p))$$

where $CoefVar(\cdot)$ denotes the coefficient of variation. Analogously, we have

$$\text{Var}\,(\log(L)) \approx \frac{\text{Var}\,(L)}{\text{E}(L)^2} = CoefVar^2(L).$$

Therefore, (3) implies (4) and concludes the proof. □

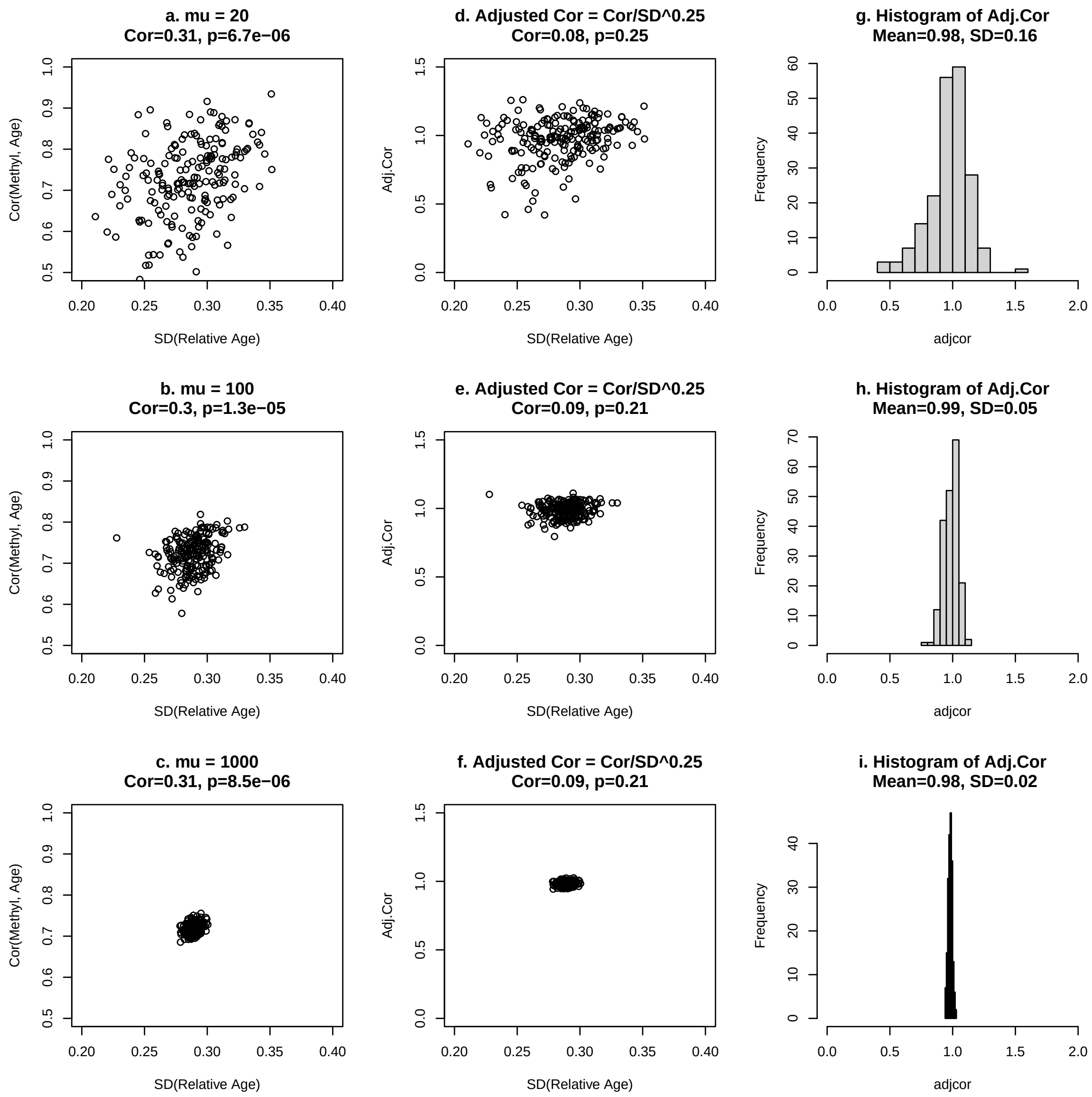


Figure S12: **Simulation Studies: Rationale for Adjusted Correlation Definition.** Panels (a,b,c) show Age correlation cor(M,A) (y-axis) against the standard deviation of relative age (x-axis), revealing a positive Pearson correlation ($Cor \approx 0.3$) due to stratum size variation. Panels (d,e,f) depict Adjusted age correlation (y-axis) against the standard deviation of relative age (x-axis), where the diminished, non-significant Pearson correlation ($Cor \approx 0.09$) demonstrates robustness against variable animal counts. Panels (g,h,i) show histograms of adjusted age correlations, indicating the adjusted correlation's mean value is near 1, and its standard deviation decreases with sample size. These results suggest $Adj.$Cor approaches 1 as stratum size increases. Supplementary Methods provide additional details.

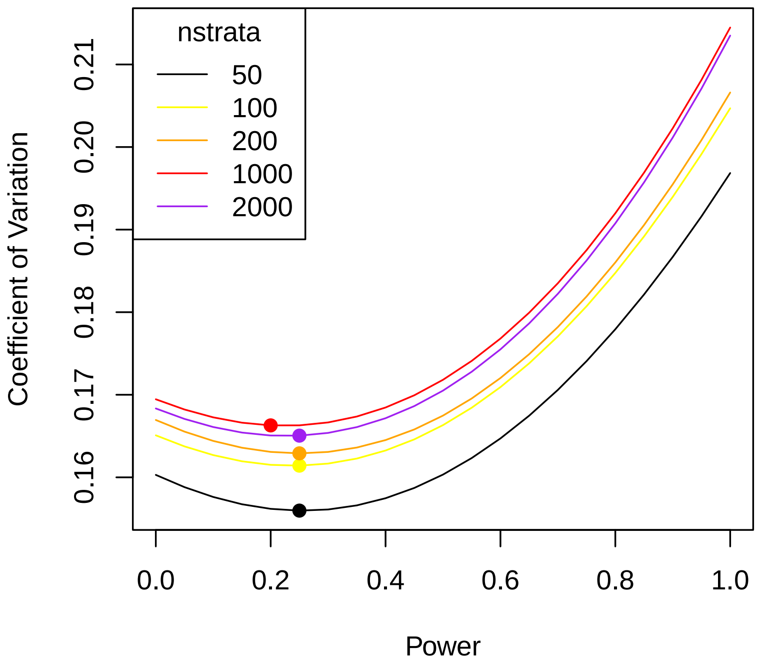


Figure S13: **Simulation studies to justify the definition of an adjusted correlation.** Supplementary Figure S12. Coefficient of Variation (CV) of Adjusted Correlations as a function of different powers. The CV displays a U-shape when the power increases, hence a minimum is achievable. The optimal power is achieved at 0.25 for most cases.

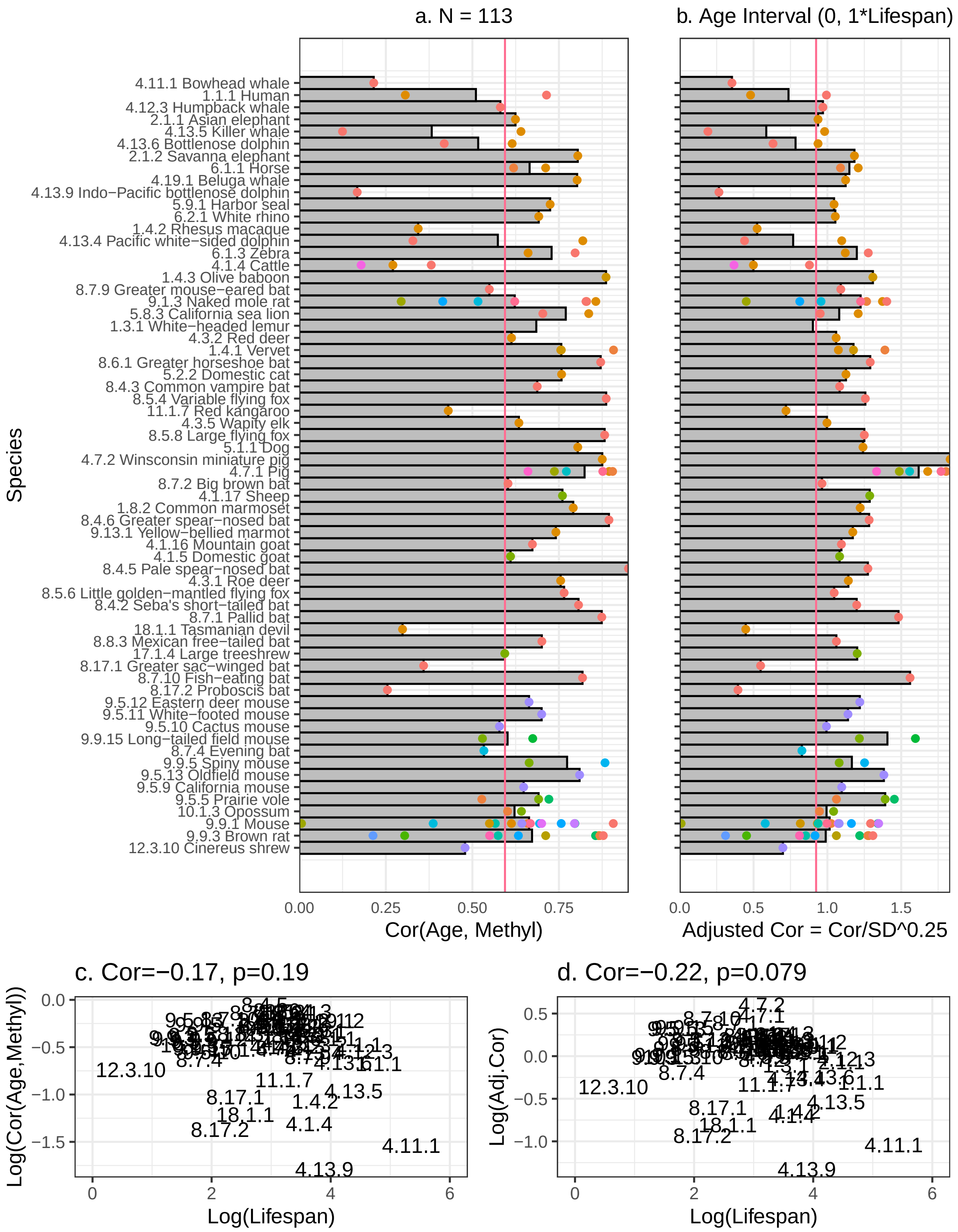


Figure S14: **Species level analysis shows that age correlations do not relate to maximum lifespan.** Barplot for $\mathrm{Cor}(Age, Methyl)$ and $Adj.\mathrm{Cor} = \frac{\mathrm{Cor}(Age,Methyl)}{\mathrm{SD}\,(\mathbf{R})^{0.25}}$ in different species (113 species tissue strata with at least 15 samples). Each horizontal bar (y-axis) corresponds to a different species. Species are sorted by maximum lifespan: from long lived (top) to short lived (bottom). Each bar length reports correlation value for each species across all tissue types, while the dots report the values for different tissue types. The QCOD for $\mathrm{Cor}(Age, Methyl)$ is 0.189 and that for $Adj.\mathrm{Cor}$ is 0.137. Mean and Median $\mathrm{Cor}(Age, Methyl)$ across species are 0.66 and 0.69 respectively, and those for $Adj.\mathrm{Cor}$ are 1.04 and 1.12. Following equation (1), Methyl was defined as mean methylation value across all CpGs located in bivalent promoter state 2 (BivProm2+). The correlation is **c.** $-0.166(p = 0.19)$ for $\mathrm{Cor}(Age, Methyl)$ and **d.** $-0.22(p = 0.08)$ for $Adj.\mathrm{Cor}$.

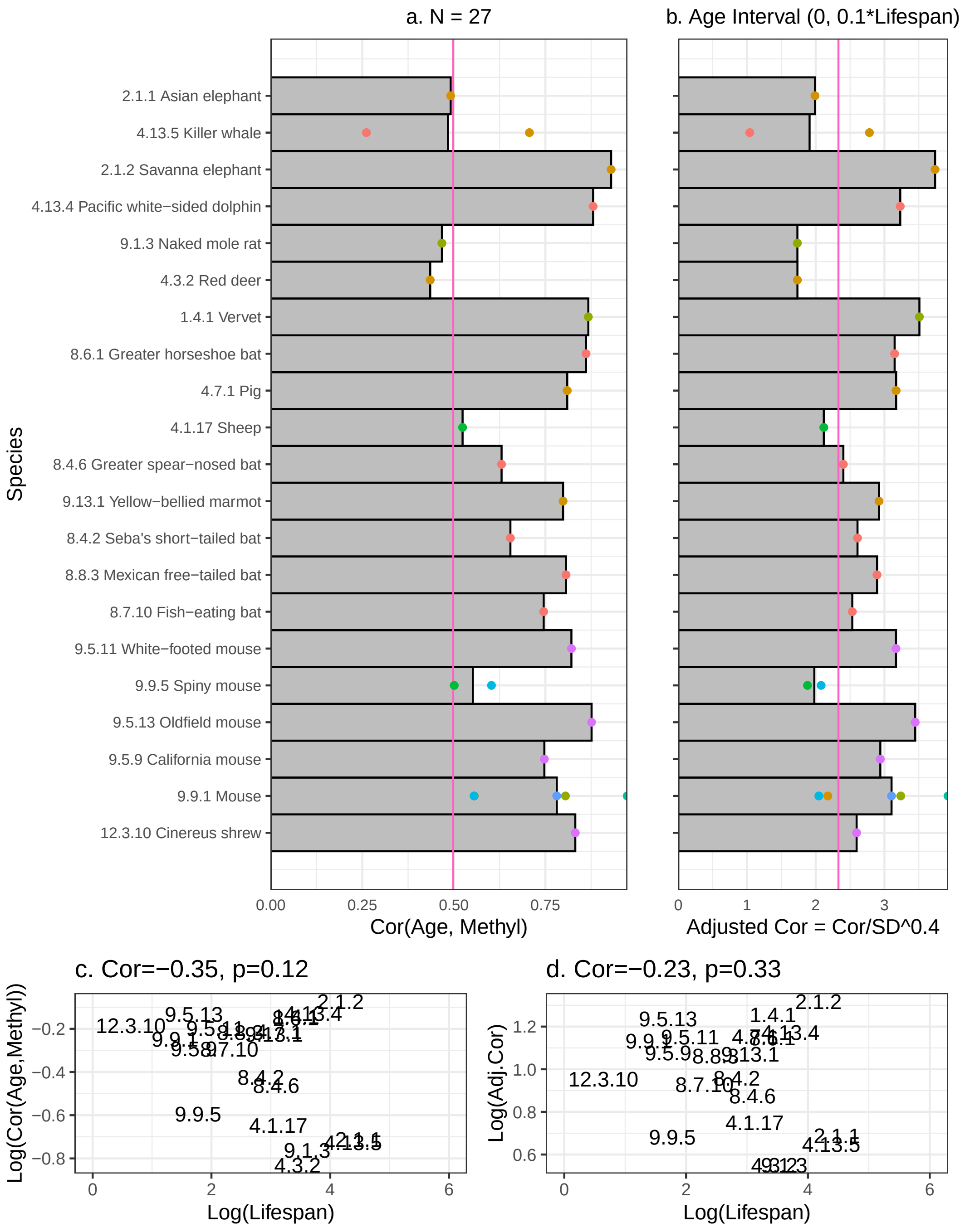


Figure S15: **Species level age correlations in young animals**. This analysis, akin to Figure S14, focuses on younger animals with relative age $R < 0.1$, and species are arranged by lifespan. Each dot represents a different tissue. Panel (a) displays Cor(Age, Mean Methylation in BivProm2+) for strata with sample size >= 15; panel (b) shows Adj.Cor for the same strata. The Mean, Median, and QCOD for Cor(Age,Methyl) are 0.5, 0.6, and 0.137 respectively; for Adj.Cor, they are 2.3, 2.6, and 0.115.

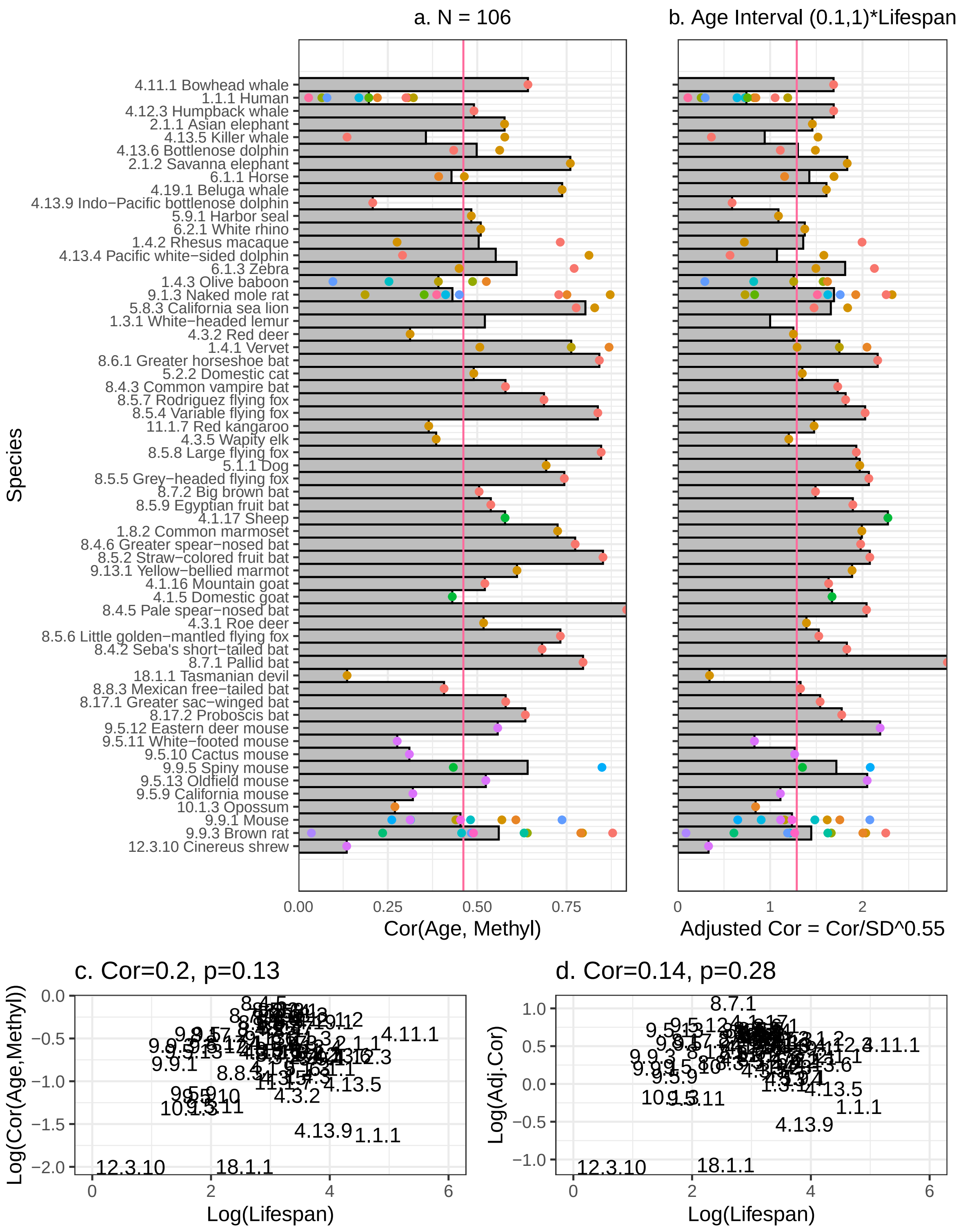


Figure S16: **Species level age correlations in old animals**. This analysis mirrors Figure S14, focusing on older animals with relative age $R \geq 0.1$, and species are sorted by lifespan. Each dot signifies a different tissue. The left panel shows Cor(Age, Mean Methylation) in BivProm2+ for strata with sample size >= 15; the right panel shows Adj.Cor for the same strata. The Mean, Median, and QCOD for Cor(Age,Methyl) are 0.46, 0.51, and 0.389 respectively, and for Adj.Cor, they are 1.17, 1.39, and 0.253.
